# Immunogenic tumor mass dormancy as a driver of persistent residual lesions in immunotherapy-treated melanoma

**DOI:** 10.64898/2025.12.11.692633

**Authors:** Yingxiao Shi, Zoltan Maliga, Tuulia Vallius, Shishir M Pant, Roxanne Pelletier, Brigette Kobs, Priyanka Solanky, Yiwen He, Eliezer Van Allen, Sandro Santagata, Patrick Ott, Christine G Lian, Elizabeth I. Buchbinder, David Liu, Peter K Sorger

**Affiliations:** Laboratory of Systems Pharmacology, Harvard Program in Therapeutic Science, Harvard Medical School, 200 Longwood Avenue, Boston, MA 02115; Ludwig Center at Harvard; Division of Population Sciences, Dana-Farber Cancer Institute, 450 Brookline Avenue, Boston, MA 02215; Department of Immunology, Harvard Medical School; Parker Institute for Cancer Immunotherapy, Dana-Farber Cancer Institute, Boston, MA; Department of Pathology, Brigham and Women’s Hospital, Harvard Medical School, 75 Francis Street, Boston, MA 02115; Department of Dermatology, Brigham and Women’s Hospital, Harvard Medical School, 75 Francis Street, Boston, MA 02115; Department of Dermatology, Massachusetts General Hospital; Department of Medical Oncology, Dana-Farber Cancer Institute, 450 Brookline Avenue, Boston, MA 02215; Department of Systems Biology, Harvard Medical School, 200 Longwood Avenue, Boston, MA 02115

## Abstract

Tumor dormancy is thought to enable cancer recurrence, but the settings and mechanisms of dormancy in patients are poorly characterized. Both immunogenic tumor mass dormancy, involving continued tumor cell proliferation and balanced immune-mediated cell death, and cell-level quiescence have been observed in preclinical models. Here, we demonstrate the existence of mass dormancy rather than quiescence in long-term stable residual lesions in melanoma patients treated with immune checkpoint inhibitors (ICI). In a large stable lesion subjected to detailed spatial profiling, the proportion of proliferating tumor cells is similar to that in site-matched tumors from patients progressing after ICI. Residual stable lesions from other patients, judged to contain only scar tissue upon clinical pathology review, also contained nests of dividing tumor cells surrounded by active immune cells. These findings demonstrate the presence of viable tumor cells and mass dormancy in persistent stable lesions, a finding with implications for disease monitoring and management.

## INTRODUCTION

Tumor dormancy, tumor cell quiescence, and therapy-resistant residual disease are interconnected phenomena thought to play important roles in cancer relapse, metastasis, and long-term survival. Dormancy at a single cell level has attracted the greatest recent interest and involves tumor cells that enter a prolonged period of quiescence or senescence with little evidence of proliferation.^1,2^ Tumor “mass dormancy” is an alternative form of dormancy in which cancer cells continue to proliferate, but tumors remain constant in size due to a balance between proliferation and cell death^3^. In “angiogenic” tumor mass dormancy, cancer cells proliferate but tumor mass is controlled by programmed cell death (PCD) induced by metabolic and growth limitations.^4^ A similar phenomenon has been observed in cultured tumor cells exposed to anti-cancer drugs: what appears to be a population-level cytostasis (no net growth due to cell cycle arrest) is often a dynamic state with matched (or nearly matched) division and death rates.^5^ Immunogenic tumor mass dormancy^6^ is a related phenomenon in which surveillance by the adaptive immune system (T cells in particular) kills proliferating tumor cells in rough balance to the numbers of cells that are born. These mechanisms of dormancy have been demonstrated in preclinical models,^7^ and a transient dormant state has been postulated to occur during immunoediting prior to tumor cell escape,^8^ but the forms of dormancy occurring in human tumors, and the clinical settings in which they might occur, have not been well demonstrated. In particular, immunogenic tumor mass dormancy has not, to our knowledge, been demonstrated as a mechanism of tumor dormancy in patients following immunotherapy.

We therefore investigated residual lesions in melanoma patients treated with immune checkpoint inhibitors (ICIs). ICIs have revolutionized the treatment of metastatic melanoma, with long-term studies documenting ten-year survival rates as high as 44-52%.^9,10^ In a substantial subset of responsive patients (anecdotally ∼20-40%)^11^ tumors shrink on therapy but remain palpable or visible by radiological imaging for an extended period of time without evidence of growth or metastasis.^12^ The clinical and biological interpretation of residual lesions is unclear, and they are not routinely removed in the absence of other evidence of disease progression. However, a recent clinical study by Buchbinder et al.^13^ was successful in obtaining biopsy or resection specimens from persistent stable lesions at least 2 years post-ICI initiation, providing a unique resource for studying mechanisms of tumor dormancy post-ICI treatment. In this study, we perform detailed multi-modal spatial profiling to identify the underlying mechanisms of tumor persistence and dormancy.

One large lymph node metastasis in the cohort (specimen MEL101) was stable on CT imaging for 35 months but found to be FDG-avid on PET/CT and was biopsied on study. It was subsequently resected based on evidence of viable tumor cells in the study biopsy. The resulting specimen is unique in being large enough for detailed analysis by spatial proteomics, spatial transcriptomics, and single cell RNA sequencing (scRNA-seq). Analysis of MEL101 revealed the co-occurrence of a high fraction of proliferating tumor cells, active T-cell mediated immune editing, expression of quiescence markers, and several types of PCD. The frequency of these states varied across the tumor, but in no case was quiescence the dominant phenotype. Moreover, none of the cell states found in this tumor (e.g., dedifferentiated, neural crest-like, or melanocytic) were enriched in quiescent cells. We conclude that this exceptional post-ICI tumor is characterized by one or more forms of tumor mass dormancy rather than true quiescence. We also identified viable and proliferative tumor cells in another patient with a pathologically negative tumor (clinically evaluated by a dermatopathologist, based on H&E images, to be disease-free with scar-tissue only), along with surrounding T cells and other immune cells. Thus, immunogenic tumor mass dormancy rather than quiescence likely underlies clinical non-progression in two of six post-ICI residual lesions examined. This suggests mass dormancy may arise at substantial frequency in this clinical setting.

## RESULTS

The Buchbinder et al.^13^ melanoma cohort used in this study comprises specimens from metastatic melanoma patients treated with ICIs at least 2 years prior to specimen collection (biopsy or resection). These patients were treated with at least two doses of ipilimumab, nivolumab, or ipilimumab plus nivolumab and then followed by CT or PET/CT imaging at regular intervals. Tumors were scored as exhibiting a long-term stable state based on the absence of progressive increases in radiological signal for ≥ 6 months (see **Supplementary Table 1** for clinical metadata). Biopsy or resection specimens of these lesions were available for six of these patients (MEL101 to 106), with their clinical course and disease characteristics summarized in **Fig. 1a**. As previously reported, only one of the six sampled lesions (MEL101), a lymph node metastasis, contained viable tumor cells detected by pathologist review of H&E-stained images. The other samples contained melanophages (macrophages that have ingested melanin) and necrotic or scar tissue but were scored as having no viable tumor cells. To compare persistent stable lesions to progressive metastatic melanoma, we acquired biopsies from three patients exhibiting progressive lymph node disease after ICI therapy (MEL107–109; **Fig. 1a**; **Supplementary Fig. 1a, b**). Freshly cut FFPE sections from these specimens were H&E-stained and subjected to expert pathology review to confirm tumor involvement (**Fig. 1a**). Adjacent sections were characterized by multiple spatial profiling methods, including cyclic immunofluorescence (CyCIF) imaging,^14^ GeoMx microregional spatial transcriptomics. CyCIF was performed with multiple antibody panels to enable discrimination of tumor differentiation states (e.g., SOX10, MART1, PMEL, and NGFR), proliferation (KI67, PCNA, and phospho-histone H3 (pH3)), apoptosis (cleaved caspase 3 (cCC3) and cPARP1), stromal structures (CD31 and αSMA), as well as immune lineages (e.g., CD45, CD3E, CD8A, CD4, FOXP3, CD20, CD11C, CD68, and CD163), and their activity states (e.g., PD1 and GZMB). Panels, abbreviations, and key properties of these markers are described in **Supplementary Fig. 2a, b** and **Supplementary Table 2, 3**.

**Figure 1.**
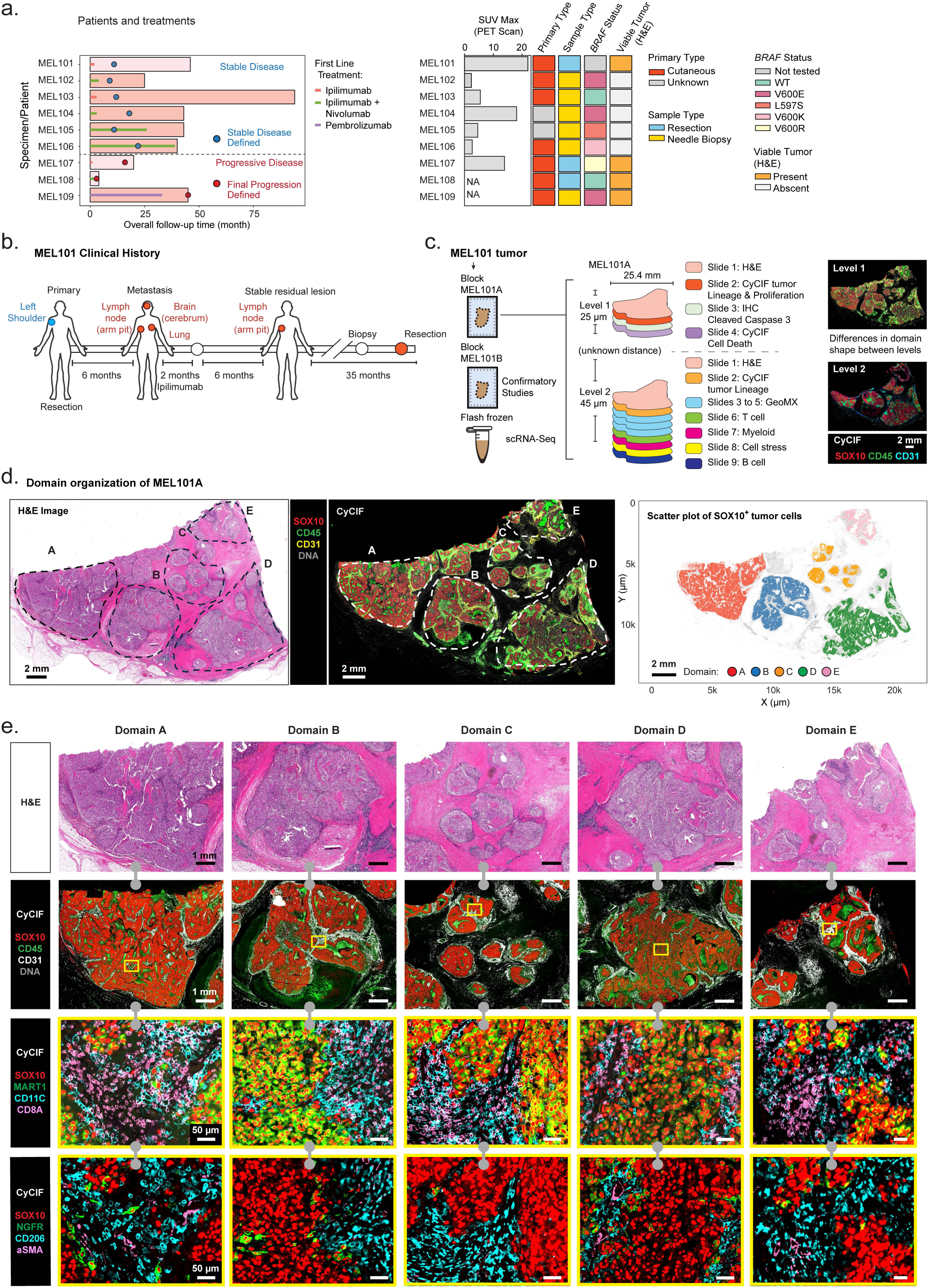
Multimodal profiling of patient MEL101 and the study cohort. **a.** Overview of the patient cohort summarizing the treatment history and clinical features of patients included in this study. **b.** Clinical timeline of patient MEL101. Patient MEL101 was initially diagnosed with a primary melanoma on the left shoulder, which was surgically resected. Six months later, metastatic lesions were detected in the axillary lymph node, cerebrum, and lung. The patient subsequently received two months of standard-dose ipilimumab treatment, which resulted in regression of all metastatic lesions except for the lymph node metastasis. This lesion was classified as a stable residual lesion six months after treatment. Serial ^18^FDG-PET/CT scans were performed every 3–6 months, and the residual lesion was surgically resected 35 months later. **c.** Schematic of tumor specimens and multimodal experimental approach. The MEL101 tumor yielded multiple tumor blocks and a frozen specimen used for single cell RNASeq. FFPE block MEL101A was used to generate multiple 5-micron sections in which Levels 1 and 2 refer to different regions of the same tumor separated along the Z-axis by an unknown amount of tumor removed from the block by other investigators. The tumor differed slightly in morphology from Levels 1 and 2, but the key features and five tumor domains were preserved. Block MEL101B was used in confirmatory studies to check overall tumor architecture. **d.** Overview of histopathological domains in specimen MEL101A. Domains A-E are indicated with dashed contours. *Left*: H&E-stained section; *Middle*: CyCIF image showing SOX10 (red), CD45 (green), CD31 (yellow), and DNA (grey); *Right*: scatter plot of SOX10^+^ tumor cells annotated by tumor domains. Scale bars, 2 mm. **e.** Magnified regions of annotated tumor domain A–E; grey barbell icons indicate that images are from the same or serial sections. *Top*: H&E-stained fields of view annotated with domain labels. *Bottom*: CyCIF images of adjacent sections showing staining for a variety of tumor and immune markers. *Row 2*: SOX10 (red), CD45 (green), CD31 (white), and DNA (grey). *Row 3*: SOX10 (red), MART1 (green), CD11C (cyan), and CD8A (pink). *Row 4*: SOX10 (red), NGFR (green), CD206 (cyan), and αSMA (pink). The regions outlined with yellow rectangles in *Row 2* represent the magnified regions shown in *Rows 3* and *4*. See **Supplementary Table S3** for how markers were used to identify different cell types. Scale bars, 1 mm and 50 µm.

MEL101, the largest specimen, was subjected to the most detailed analysis; this lymph node metastasis was obtained from a patient who had a primary melanoma on the left shoulder that was treated with wide local excision. Six months later, the patient presented with metastases in the left axillary lymph node, brain, and lungs (**Fig. 1b**). This patient received four cycles of ipilimumab (anti-CTLA4) over two months, with an initial response observed in all lesions. Ultimately, all lesions regressed except for the left axillary lymph node metastasis that remained radiographically stable on CT for 35 months (**Fig. 1b**). Biopsy and H&E imaging revealed the presence of tumor cells and the lesion was therefore resected. Of the six patients with stable residual lesions biopsied, this was the only case in which tumor cells were detected via standard pathologic evaluation. Because of its size, this specimen provides a unique opportunity to perform detailed spatial profiling of a stable tumor. Fourteen 5 μm sections from two tissue blocks were therefore subjected to 30 to 64-plex CyCIF and 16-plex ORION imaging^15^ (ORION is a one-shot multiplexed fluorescence imaging method that is particularly effective with fragile biopsy specimens) and immunohistochemistry (IHC) (**Fig. 1c; Supplementary Fig. 2c** and **Supplementary Table 2, 4**). For GeoMx, 133 microregions (MRs) representing tumor and non-tumor regions were selected for characterization based on tissue morphology markers (**Supplementary Fig. 2c–e**) and CyCIF images of adjacent sections. Each MR contained ∼125–1500 cells. Prior to mRNA preparation, some MRs were separated into tumor and immune compartments (this process is referred to as “segmentation” in the GeoMx workflow, but we refer to it as “enrichment” to avoid confusion with the conventional use of segmentation in image analysis). An adjacent fresh frozen tumor tissue from block 1 was used for scRNA-seq (**Fig. 1c; Supplementary Fig. 2f, g**) and a single section from block 2 was subjected to 10x Genomics Xenium spatial transcriptomics using the Xenium 5K panel with morphology expansion markers (ATP1A1, CD45, E-cadherin, and DAPI) (**Supplementary Fig. 2h, i**).

Sections of MEL101A obtained at two levels of the tissue block (Levels 1&2) contained multiple distinct tumor and immune cell-rich domains (**Fig. 1c**), each of which was 1–8 mm in greatest dimension (∼ 1–14 × 10^4^ cells each in 2D sections) separated by bands of fibrosis. Based on this overall structure we grouped the tumor into 5 separate domains (A–E; dashed lines in **Fig. 1d**). The proportions of tumor and immune cells and the extent of necrosis and fibrosis varied among the five domains with A being the most cellular and E the most fibrotic (**Supplementary Fig. 3a–b**). Each of the five domains contained a mixture of tumor, stromal and immune cells (**Fig. 1e**) that were found at a microanatomical level to correspond to tumor bed, peritumoral, necrotic, and perivascular regions (**Fig. 2a, b**; see METHODS). An adjacent specimen (MEL101B) had a similar organization but with less intervening fibrotic tissue (**Supplementary Fig. 2j**). Averaged across the five tumor domains, ∼58% of cells were SOX10^+^ tumor cells whereas 42% were lymphocytes, myeloid cells, and stromal cells of various types (**Fig. 2b–d; Supplementary Fig. 3a**). Considering the specimen as a whole, the vast majority (95%) of lymphocytes and myeloid cells were found in peritumoral and perivascular regions rather than within the tumor bed itself (**Fig. 2d; Supplementary Fig. 3c, d**). Necrotic regions, which were particularly prominent within tumor Domain A, contained cellular debris of various types and frequently stained positive for the CD66B cell surface glycoprotein, which is consistent with tumor necrosis with neutrophilic response (**Fig. 2b; Supplementary Fig. 3b –** Necrosis).

**Figure 2.**
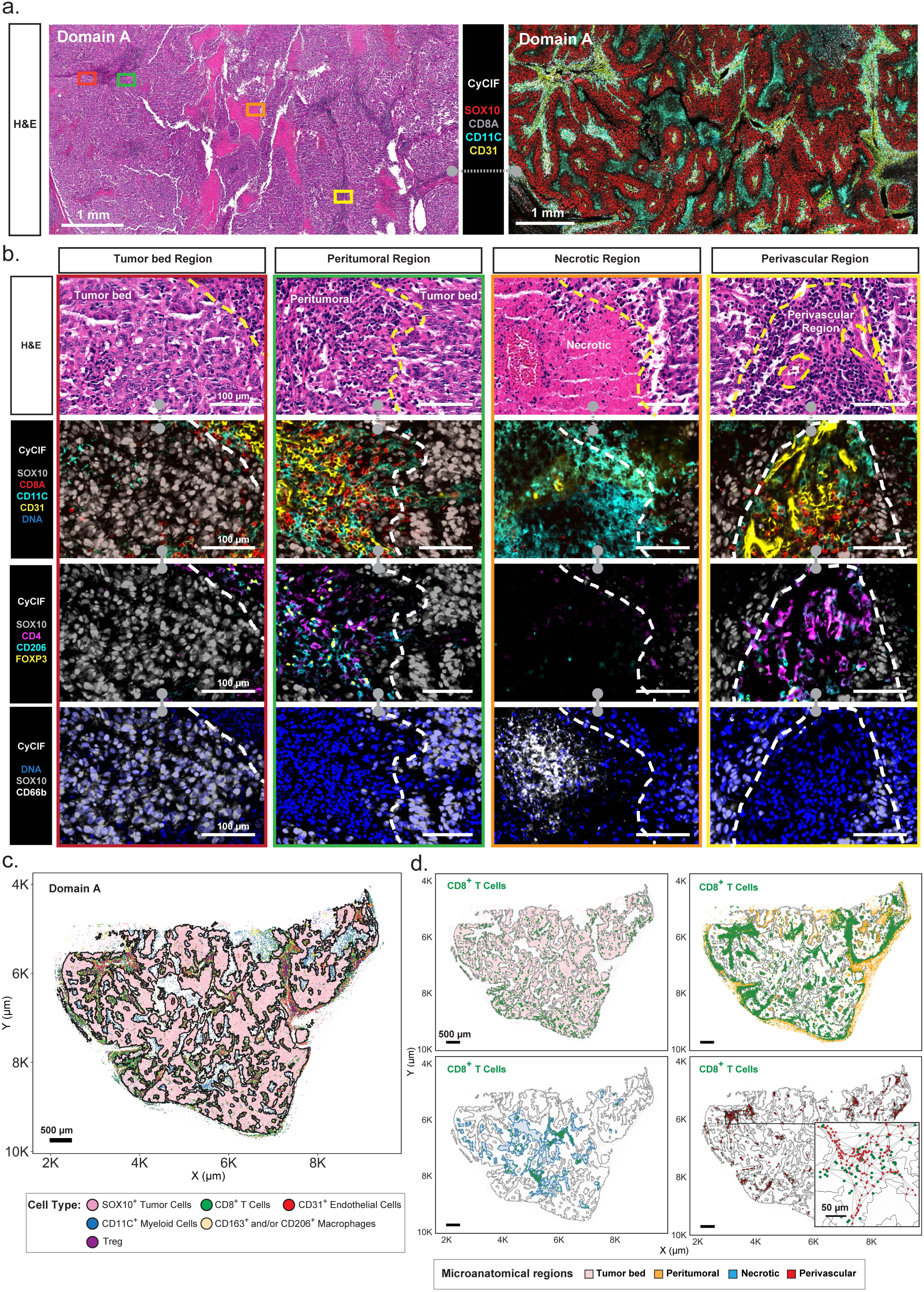
Characterization of MEL101 histopathological domains and microanatomical regions. For H&E-stained and CyCIF images, the grey barbell icon denotes the same CyCIF image with different channels shown; the barbell with dashed connector denotes adjacent sections. **a**. Field of view of MEL101 Domain A (sections 1 and 2 from MEL101A Level 1) showing H&E-stained image (left) and corresponding CyCIF image (right) with selected markers of a 64-plex image shown: SOX10 (red), CD8A (grey), CD11C (cyan), and CD31 (yellow). The regions outlined with rectangles represent the magnified regions shown in panel *b*. The color of each rectangle denotes a different microanatomical region: Tumor bed (red), Peritumoral (green), Necrotic (orange), and Perivascular (yellow). Scale bars, 1 mm. **b.** Magnification views of four representative microanatomical regions in Domain A (as depicted by colored boxes in panel *a*). From left to right: Tumor bed region, Peritumoral region, Necrotic region, and Perivascular region*. Row 1*: H&E-stained microanatomical regions. *Row 2* (CyCIF): SOX10 (grey), CD8A (red), CD11C (cyan), CD31 (yellow), and DNA (blue). *Row 3* (CyCIF): SOX10 (grey), CD4 (purple), CD206 (cyan), and FOXP3 (yellow). *Row 4* (CyCIF): SOX10 (grey), CD66 (white), and DNA (blue). Scale bars, 100 µm. The yellow and white dashed lines denote tumor boundary defined by SOX10^+^ tumor cells. **c.** Scatter plot displaying the spatial distribution of selected cell types within Domain A. The black outline denotes tumor boundary defined based on SOX10^+^ tumor cells. Each dot represents a single cell. Scale bar, 500 µm. **d.** Scatter plots showing the spatial distribution of CD8^+^ T cells (green) within the microanatomical regions: tumor bed (pink, top left), peritumoral (orange, top right), necrotic (blue, bottom left) and perivascular (red, bottom right) regions. The black rectangle indicates a magnified perivascular region with CD8^+^ T cells (green dots) and CD31^+^ cells (red dots). Black lines in the perivascular regions represent a Delaunay-based connectivity network between CD31^+^ cells. Scale bars, 500 µm and 50 µm.

Pathology review of H&E data revealed the presence of multiple features of immune-related tumor regression^16^ including tumor necrosis, stromal fibrosis, and immune infiltration into the tumor bed. Domain A had the greatest level of active tumor killing and necrosis with neutrophilic and lymphocytic infiltration; Domain B exhibited immune infiltration with the emergence of early stromal fibrosis; Domain C and D, were characterized by prominent stromal fibrosis, a later stage of immune regression, and Domain E was largely fibrotic. We hypothesized that MEL101 Domains A–E correspond to different stages of the regression process and sought to confirm this using CyCIF and GeoMx data.

### Tumor cell proliferation in persistent stable and progressing lesions

Analysis of CyCIF images showed that ∼12% of tumor cells in MEL101 were positive for the KI67 proliferation marker (as were 35% of immune cells; **Fig. 3a, b**; **Supplementary Fig. 4a**); unexpectedly, this was similar to the average proportion of KI67^+^ tumor cells in progressing lymph node tumors in ICI-treated patients (MEL107–109; range 3% to 12% in **Fig. 3a, b**; **Supplementary Fig. 4b**). scRNA-seq data provided further evidence that cells were actively dividing in MEL101: a tumor cell cluster comprising ∼5.4% of sequenced cells was enriched in G1/S, G2/M (both *P* < 0.0001), and other related signatures (cluster 4; **Fig. 3c–e**). We also analyzed tissue using a CyCIF panel that included markers specific to multiple stages of the cell cycle. The approximate order of expression (or appearance) of these markers in freely proliferating cells is as follows: the p21 and p27 cyclin dependent kinase (CDK) inhibitors are specific to cell cycle arrest and exit (p27 is also regarded as a tumor quiescence marker)^17^, Cyclin D1 (CCND1) to G1/S cells, Cyclin E1 (CCNE1) and the DNA licensing factors PCNA and Geminin to S phase cells, Cyclin A2 (CCNA2) and Cyclin B1 (CCNB1) to G2/M cells, and phospho-histone H3 (pH3) to mitotic cells (**Fig. 3f**; **Supplementary Fig. 4c**). We found that adjacent tumor cells expressed these markers in multiple combinations consistent with their occupying different phases of the cell cycle, as expected for proliferating cells.

**Figure 3.**
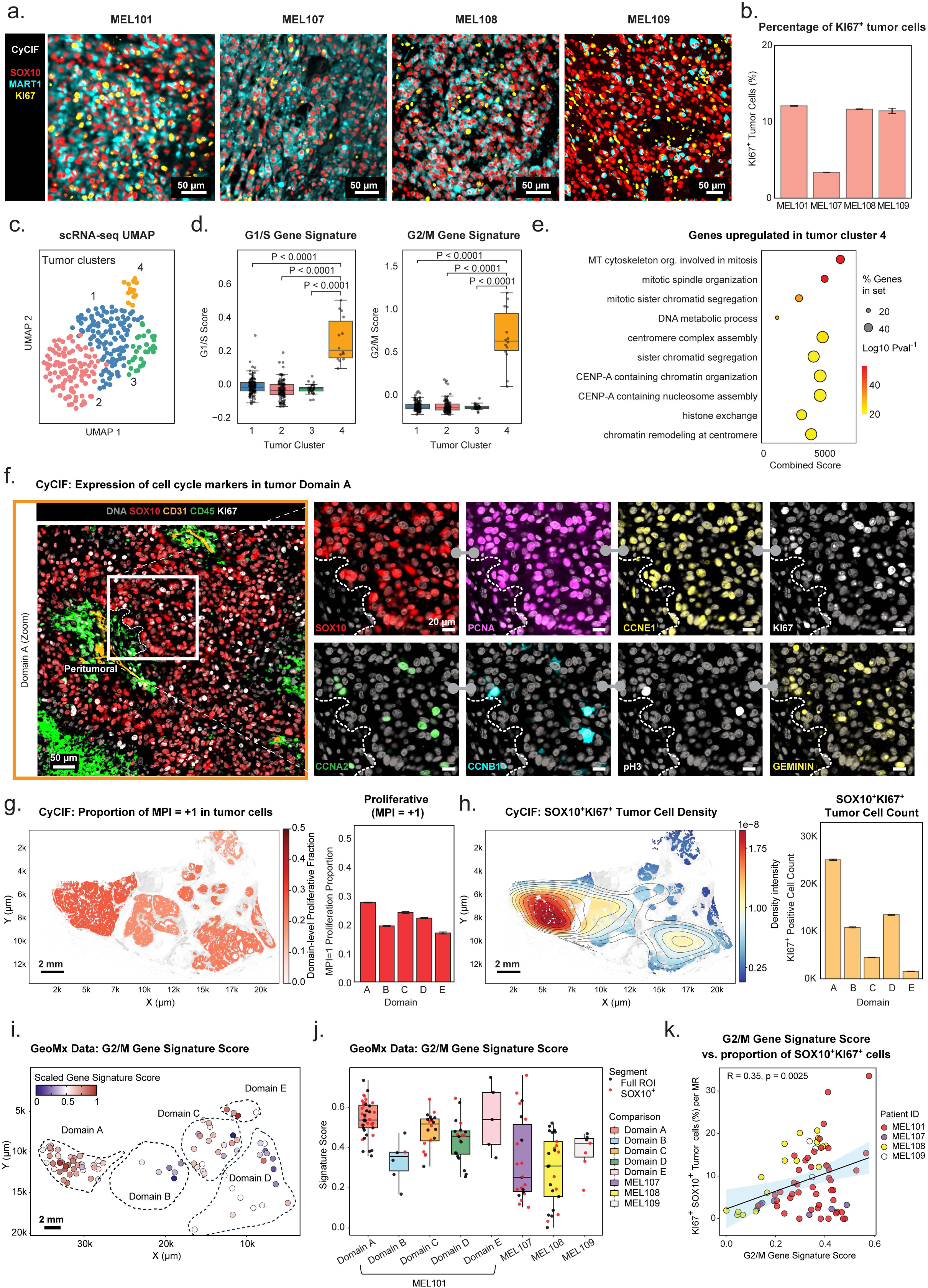
Tumor cell proliferation in persistent stable and progressing lesions. **a.** Magnified CyCIF images showing selected channels from a 42-plex CyCIF image: SOX10 (red), MART1 (cyan), and KI67 (yellow) within MEL101 and progressing tumors (MEL107-MEL109). Scale bars, 50 µm. **b.** Bar plot showing the percentage of SOX10^+^KI67^+^ tumor cells relative to all tumor cells in specimens MEL101 and MEL107–MEL109. Error bars represent binomial standard deviations. **c.** UMAP of tumor cells (scRNA-seq data) colored by tumor cell cluster ID. The clusters were obtained by unsupervised Leiden clustering. **d.** Boxplots comparing G1/S (left) and G/2M (right) gene signature scores across tumor clusters 1–4 from panel *c*. Boxes represent the first and third interquartile of the data; whiskers extend to show the rest of the distribution except for points identified as outliers. Statistical significance was assessed using the Mann–Whitney U test. **e.** Dot plot representing pathway enrichment analysis of genes upregulated in tumor cluster 4. Dot size reflects the proportion of enriched genes in each gene set, while color indicates statistical significance (−log₁₀ *P*-value). The combined score represents the strength of enrichment for each pathway**. f.** Selected channels of a 64-plex CyCIF image from domain A of MEL101 showing cell cycle markers. *Left panel*: staining for DNA (grey), SOX10 (red), CD31 (orange), CD45 (green), and KI67 (white) within Domain A. The region outlined with a white rectangle represents the magnified region shown in the right panel; *Right panels*: magnified field of view stained for SOX10 (red), PCNA (purple), CCNE1 (yellow), KI67 (white), CCNA2 (green), CCNB1 (cyan), pH3 (white), and GEMININ (yellow). Grey barbell icon denotes the same CyCIF image with different channels shown. The white dashed lines denote tumor boundary defined by SOX10^+^ tumor cells. Scale bars, 50 µm and 20 µm. **g.** Spatial distribution and quantification of proliferative tumor cells (MPI = +1). *Left*: scatter plot of SOX10^+^ tumor cells in specimen MEL101A. Cells are color-coded by the average proportion of proliferative cells within each domain. Scale bar, 2 mm*. Right*: bar plot showing the proportion of MPI= +1 cells within each domain. Error bars represent binomial standard deviations. **h.** Spatial distribution and quantification of SOX10⁺ KI67⁺ proliferative tumor cells. *Left*: Kernel density estimation (KDE) plot highlighting the spatial density of SOX10⁺ KI67⁺ tumor cells across entire tumor. A Gaussian KDE was applied to the centroid coordinates of double-positive cells, with intensity clipped at the 98^th^ percentile and regions below the 10^th^ percentile masked for contrast enhancement. Contour lines denote high-density regions, overlaid on all cells (grey). The color scale represents kernel density-estimated spatial intensity (range: 0–1.8 × 10^−8^, scaled unit) *Right*: The absolute abundance of SOX10^+^KI67^+^ tumor cells across different domains. Scale bar, 2 mm. **i.** Scatter plot showing spatially resolved G2/M gene signature scores across GeoMx tumor MRs. Each data point represents one tumor MR, and color indicates the scaled gene signature score. Scale bar, 2mm. **j.** Boxplot comparing G2/M gene signature scores across tumor GeoMx MRs in Domains A–E and progressing tumors (MEL107–MEL109). Each dot represents a tumor MR (either SOX10⁺ segmented as part of the GeoMx RNA protocol or full MRs). Boxes represent the first and third interquartile of the data; whiskers extend to show the rest of the distribution except for points identified as outliers. **k**. Scatter plot showing the Pearson correlation between the G2/M gene signature scores (x-axis, GeoMx) and the percentage of SOX10^+^KI67⁺ tumor cells among all tumor cells (y-axis, CyCIF). Each data point represents an individual full tumor MR colored by patient ID. The solid line indicates the linear regression fit, with a 95% confidence interval shown (shaded area). Significance was calculated using a two-sided Pearson correlation test.

To integrate cell cycle data into a single multi-marker metric, we calculated the Multivariate Proliferation Index (MPI)^18^ (**Fig. 3g; Supplementary Fig. 4d, e**). This confirmed the presence of proliferating tumor cells (MPI = +1) in all five tumor domains in roughly similar proportions (20–28% of tumor cells); GeoMx data also confirmed the enrichment of proliferation gene signatures in tumor MRs from all five domains. The greatest density of proliferating tumor cells (cells per unit area) was observed in Domain A (**Fig. 3h**). GeoMx gene signature scores were also higher in this than other MEL101 domains or in MEL107 to 109 (**Fig. 3i, j**; **Supplementary Fig. 4f, g**). This is expected since gene signature scores for an MR vary with the density of signature-positive cells within that MR, whereas the MPI proliferation index measures division on a per-cell basis. In sum, CyCIF, scRNA-seq, and GeoMx data robustly and consistently demonstrated proliferation of tumor cells in MEL101, a clinically stable lesion, at levels comparable to tumor cell proliferation in lesions progressing on ICI therapy (**Fig. 3k; Supplementary Fig. 4h**).

### Immune surveillance of tumor domains in MEL101

The proportion of CD45^+^ immune cells in MEL101 tumor domains varied from 12 to 40%, demonstrating extensive infiltration (**Supplementary Fig. 3a**). Hierarchical cell type calling of CyCIF data identified seven CD8^+^ T cell states: (1) naïve T cells that co-expressed LEF1 and CD45RA, (2) memory T cells that expressed CD45RO, (3) resident memory T cells that co-expressed CD45RO and CD103, (4) cytotoxic T cells (CTLs) that expressed the cytolytic serine protease granzyme B (GZMB) as well as the PD1, LAG3, and TIM3 checkpoint proteins at intermediate levels, (5) CTLs similar to (4) but also expressing KI67, CXCL13, and pLCK (Tyr 394), a tyrosine kinase associated with the T cell receptor (TCR), (6) partially exhausted T cells that co-expressed PD1 and LAG3 with little or no TIM3, and (7) terminally exhausted T cells that co-expressed PD1, LAG3, and TIM3 (**Fig. 4a, b; Supplementary Fig. 5a**). The proportions of these cell types (relative to immune cells as a whole) differed by ∼200-fold, with CTLs (Cytotoxic and Proliferative Cytotoxic, n = 21.5k) and memory T cells (Memory and Resident Memory, n = 23.1k) the most common and naïve T cells the least (cluster 1; n = 71). Naïve T cells were primarily localized to one organized lymphoid structure while other T cell subtypes were distributed across the tumor domains (**Supplementary Fig. 5d–f**; **Fig. 4e–g**).

**Figure 4.**
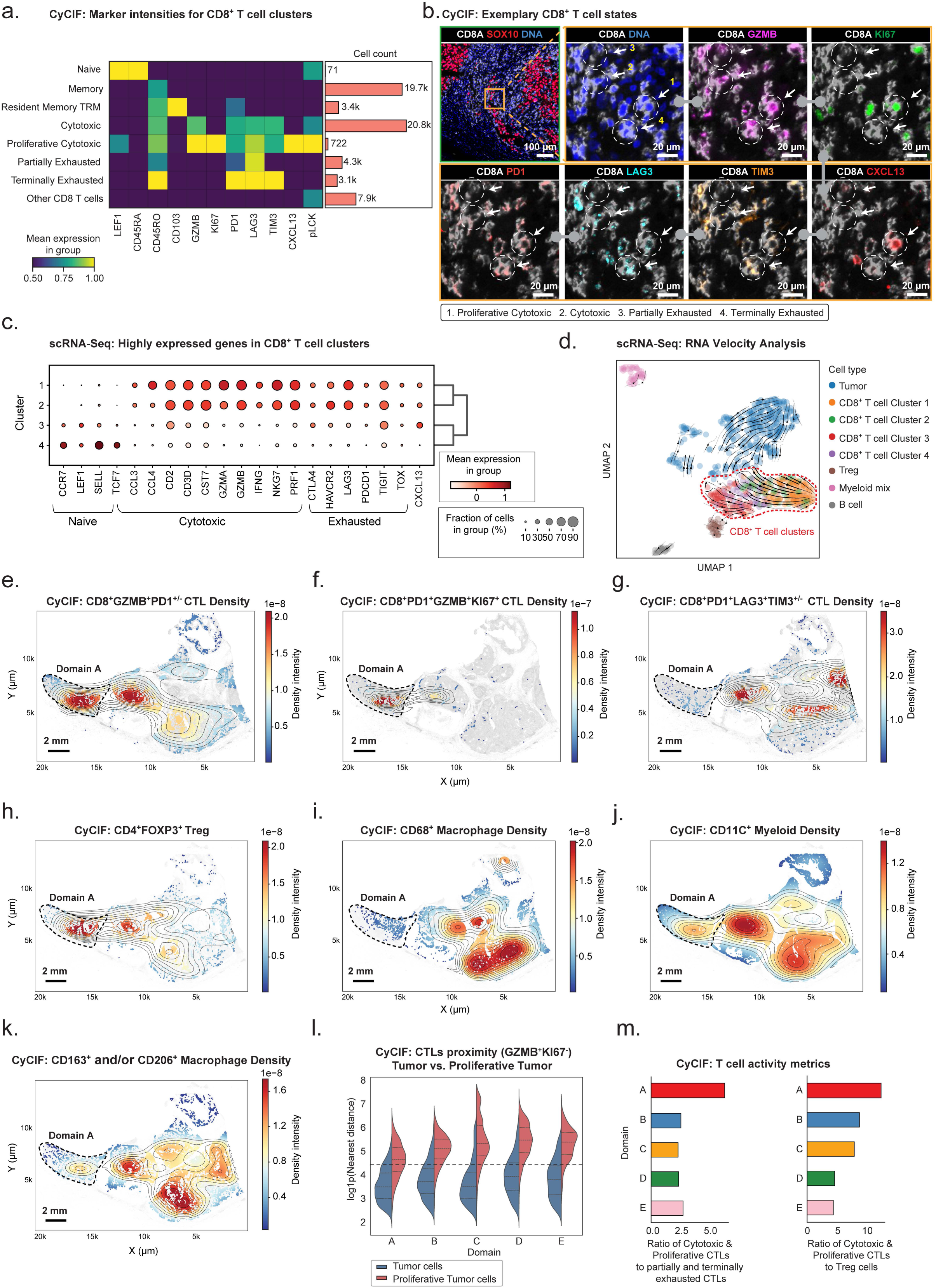
Immune surveillance of tumor domains in MEL101. **a.** Heatmap showing the mean scaled expression of selected CyCIF marker intensities across CD8⁺ T cell subtypes with assignment of their likely states: naïve, memory, tissue-resident memory, cytotoxic, proliferative cytotoxic, partially exhausted, terminally exhausted, and other. Bar plots on the right denote the absolute cell count in each subtype. **b.** CyCIF images of representative CD8^+^ T cell subtypes in a 41-plex CyCIF image showing staining for CD8A (white), SOX10 (red), DNA (blue), GZMB (magenta), KI67 (green), PD1 (red), LAG3 (cyan), TIM3 (orange), and CXCL13 (red). The white dashed circles and numbers highlight CD8^+^ T cell subtypes. Scale bars, 100 µm and 20 µm. **c.** Dot plot depicting expression of representative genes among CD8⁺ T cell clusters identified by Leiden clustering in scRNA-seq data. Dot size reflects the percentage of cells expressing each gene, while color intensity represents the average normalized expression level. Marker genes are grouped by functional categories: naïve, cytotoxic, and exhausted based on probable function. **d.** UMAP of scRNA-seq data colored by cell type. RNA velocity vectors overlaid on the UMAP illustrate dynamic state transitions, with a focus on the CD8⁺ T cell compartment (highlighted with a red dashed line) **e-k.** KDE plots showing spatial distribution of specific immune cell types across MEL101 domains. Contour lines indicate regions of high local density, overlaid on all cells (grey). The color scale represents kernel density-estimated spatial intensity (range in scaled units as shown). The black dashed region outlines Domain A. Scale bars, 2 mm. **e.** CD8A^+^GZMB^+^ CTLs with PD1 status undefined. **f.** Proliferative cytotoxic CD8^+^GZMB^+^KI67^+^ CTLs cells. **g.** Partially or terminally exhausted CD8A^+^LAG3^+^ and/or TIM3^+^ CTLs. **h**. CD4^+^FOXP3^+^ Treg cells. **i.** CD68^+^ macrophages. **j.** CD11C^+^ myeloid cells. **k**. CD163^+^ and/or CD206^+^ macrophages. **l.** Split violin plots of the distribution of log-transformed nearest-neighbor distances between cytotoxic CTLs (CD8^+^GZMB⁺KI67⁻) and either non-proliferative (blue) or proliferative tumor cells (red) in each domain. **m.** Bar plots showing the ratios of cytotoxic (GZMB⁺) and/or proliferative (KI67⁺) CTLs relative to partially and/or terminally exhausted CTLs (left panel) or Tregs (right panel) across Domains A–E.

scRNA-seq analysis of *CD3E^+^CD8A^+^* cells (n = 344) confirmed CD8^+^ T cell diversity. Unsupervised Leiden clustering of scRNA-seq data generated four major CD8^+^ T cell clusters (**Fig. 4c** and **Supplementary Fig. 5b, c**): Cluster 2 cells expressed the *CCL5* chemokine, pore-forming protein perforin (*PRF1*), and *GZMB* at high levels and corresponded to CTLs; Cluster 1 cells expressed the *HLA-DR* and granzyme A (*GZMA*), and corresponded to a second population of CTLs; Cluster 4 cells expressed genes found in naïve and memory T cells such as *SELL, KLF2* and *TCF7*; and Cluster 3 cells expressed high level of *CXCL13* and the *PDE3B* phosphodiesterase and likely corresponded to the CXCL13 positive CTLs identified by CyCIF. RNA velocity analysis on these data yielded a trajectory from naive CD8^+^ T cells (Cluster 4) to *CXCL13^+^CD8^+^*T cells (Cluster 3) to cytotoxic states (Clusters 1 and 2), consistent with an active T cell response (**Fig. 4d**) involving naïve, cytotoxic, and activated/exhausted T cells in proportions consistent with ongoing tumor immunoediting.

The relative abundance of T cell subtypes varied substantially with tumor domain (**Fig. 4e–g**; **Supplementary Fig. 4d–f**). Proliferative CXCL13^+^ CTLs were enriched in Domain A (**Fig. 4f**), which also contained the highest density of proliferating tumor cells (**Fig. 3g–h**). Tregs, macrophages and myeloid cells were present across all domains (**Fig. 4h–k**) with Tregs enriched in Domain A (**Fig. 4h**); macrophages (**Fig. 4i, k**) and myeloid cells (**Fig. 4j**) enriched in Domains C–E. Proximity analysis showed that CTLs were closest to proliferative tumor cells in Domain A as compared to other domains (**Fig. 4l**). Moreover, the ratio of CTLs (and proliferative CTLs) to exhausted T cells was highest in Domain A (**Fig. 4m**). These are all features of an active anti-tumor immune response and consistent with our hypothesis that Domain A represents an earlier and more active stage of immune editing than Domain B, which has a higher proportion of exhausted T cells, or Domains C–E, which have few CTLs and more abundant macrophages and fibrosis, consistent with immune downregulation and resolution.

### Programmed cell death in tumor compartments

A tumor that is actively proliferating yet remains constant in size must also experience cell death and clearance of dead cells. Tumor mass dormancy models postulate that this can be either cell-intrinsic arising from DNA damage, metabolic stress, or intrinsic defects in tumor cells or immune-mediated. Multiple forms of programmed cell death (PCD) have been described in cell lines, model organisms, and tissues including immunogenic forms of death that generate an inflammatory environment, such as pyroptosis^19^ and necroptosis^20^ (pyroptosis is also associated with better outcomes in metastatic melanoma)^21^ and non-immunogenic PCD such as apoptosis and ferroptosis.^22^ Characterization of cell death in tissues is not trivial however, since it is necessary to catch cells midway along a process in which both the protein, RNA, and DNA are degraded and prior to engulfment of dead cells by macrophages. Moreover, PCD pathways overlap in their regulatory mechanisms and RNA signatures for PCD contain many of the same genes^23^.

To maximize the ability to detect dying cells we combined IHC, CyCIF, GeoMx and Xenium 5K panel profiling. IHC was performed using antibodies against cleaved caspase-3 (cC3), the active form of the executioner caspase in apoptotic cells; CyCIF was performed with antibodies against cleaved PARP1 (cPARP1), a cC3 substrate in caspase-dependent apoptosis. Both methods identified dying cells in tumor and peritumoral microanatomical regions (**Fig. 5a, b; Supplementary Fig. 6a, b**). The majority of these cells were non-tumor cells (e.g., CD3E^+^, CD11C^+^) particularly in peritumoral regions, consistent with the short half-lives of activated T cells, neutrophils and other immune cells (**Fig. 5b, c**). However, cPARP1-positive cancer cells were also observed (**Fig. 5c – 4^th^ row**, representing ∼1% of all SOX10^+^ cells). When GeoMx transcriptomic data were examined, we observed high signature scores for necroptosis, ferroptosis, apoptosis and pyroptosis in immune MRs as well as positive scores for one or more signatures in the majority of tumor MRs (**Fig 5d; Supplementary Fig. 6c**). One challenge with these data is that even with careful selection of MRs, cross contamination of microanatomical compartments occurs (e.g. an immune-rich perivascular region within tumor-focused MR 21; **Fig 5e**). To better discriminate PCD in cancer and immune cells we examined Xenium data, which provides near single-cell resolution. We focused on cancer cells with the highest expression of one of four PCD programs in the tumor bed, perivascular, and perinecrotic regions (**Fig. 5f, g; Supplementary Fig. 6d, e**); we discounted signals in necrotic cores because nuclei were highly degraded. Microanatomical compartments were observed to have cancer cells with high scores for all four PCD programs in roughly equal proportion and without detectable spatial patterning, justifying the decision to analyze PCD in aggregate (**Fig. 5h**). PCD positive cancer cells were found throughout the tumor domain and in the absence of immediately adjacent immune cells; this may represent cell-intrinsic death or the action of immune cells that are out-of-plane. PCD was also observed in cancer cells in and around perivascular regions, and this occurred in the presence of a high level of T cell infiltration. However, dying cancer cells were most prevalent, as proportion of all cancer cells, in the perinecrotic regions (reaching ∼10% versus ∼3% in other regions **Fig 5i**) consistent with a classic mode of tumor immune editing in which necrotic cores arise from the death of cancer cells, the immune cells involved in killing them, and other cells recruited to the resulting inflammation.^24,25^

**Figure 5.**
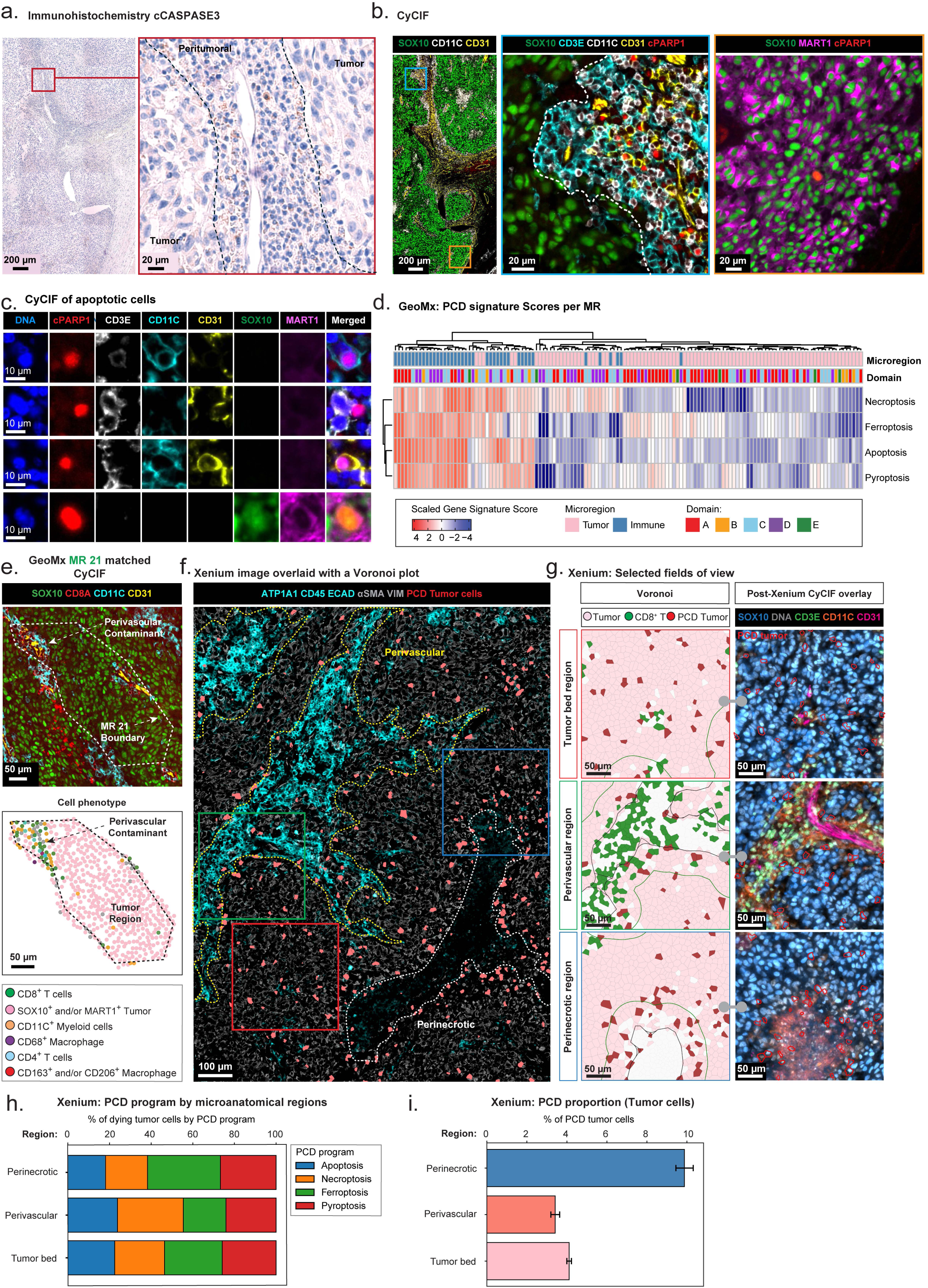
Programmed cell death in the tumor compartment in MEL101. **a.** IHC image of cleaved caspase-3 (cCASPASE3) staining within MEL101 Domain A (Level 2). The black dashed line separates the tumor and peritumoral regions. Scale bars, 200 µm and 20 µm. **b.** *Left*: selected channels from a 42-plex CyCIF showing staining for SOX10 (green), CD11C (white), CD31 (yellow). The regions outlined with blue and orange rectangles represent the magnified regions shown in the middle and right panels, respectively; *Middle*: magnified region highlighting cPARP1^+^ non-tumor cells within the peritumoral region: SOX10 (green), CD3E (cyan), CD11C (white), CD31 (yellow), and cPARP1 (red). White dashed line denotes tumor boundary defined by SOX10^+^ tumor cells; *Right*: magnified region highlighting a cPARP1^+^ tumor cell within the tumor region: SOX10 (green), MART1 (magenta), and cPARP1 (red). Scale bars: 200 µm and 20 µm. **c.** Selected channels from a 43-plex CyCIF image showing cPARP1^+^ cells: DNA (blue), cPARP1 (red), CD3E (white), CD11C (cyan), CD31 (yellow), SOX10 (green), and MART1 (magenta). Scale bar, 10 µm. **d.** Heatmap displaying gene signature scores for PCD signatures across annotated tumor Domains A–E. Columns represent individual GeoMx MRs, annotated by segment type (immune vs. tumor) and domain identity. **e.** *Top*: Selected channels of 43-plex CyCIF images (left) showing the spatial composition of tumor and immune cell subsets within tumor MR 21 with SOX10 (green), CD8 (red), CD11C (cyan); *Bottom:* Scatter plot (right) showing positions of selected tumor and immune cell types within GeoMx tumor MR 21: SOX10^+^ and/or MART1^+^ tumor cells (pink), CD8⁺ T cells (green), CD4⁺ T cells (blue), CD11C^+^ myeloid cells (orange), CD163⁺ and/or CD206⁺ macrophages (red), and CD68⁺ macrophages (purple). Scale bars, 50 µm. **f.** Representative Xenium imaging from MEL101B (Level 1) showing ATP1A1 (cyan), CD45 (cyan), ECAD (cyan), αSMA (grey), VIM (grey), and PCD tumor cells calculated based on PCD gene signature scores (PCD; red). White dashed lines denote necrotic region, and yellow dashed lines denote vascular region. The regions outlined with blue, green and red rectangles represent the magnified regions shown in *panel f*. Scale bar, 100 µm. **g.** The magnified regions of the representative tumor, perivascular, perinecrotic regions highlighted in *panel e*, illustrating the spatial distribution of PCD tumor cells across those microanatomic regions. *Left column*: Voronoi plot showing the distribution of SOX10^+^ tumor cells (pink), CD8^+^ T cells (green), and PCD tumor cells (red) within perinecrotic, perivascular and tumor regions highlighted in panel e. *Right column*: Post-Xenium CyCIF images of the corresponding regions showing SOX10 (green), CD3E (cyan), CD11C (white), CD31 (yellow). Scale bar represents 50 µm**. h.** Stacked bar plot quantifying the proportion of four different PCDs across microanatomical regions. **i.** Bar plot quantifying PCD tumor cells normalized with non-PCD tumor cells across microanatomical regions. Error bars represent the 95% confident interval around the estimated proportion of PCD tumor cells.

Considering these data as a whole, we conclude that extensive PCD is ongoing in multiple anatomical domains of MEL101 and involves a combination of immune-mediated killing of tumor cells, intrinsic tumor cell death, and death of immune cells. This is a key feature of the tumor mass dormancy model and provides the flux that opposes ongoing tumor cell birth and keeps tumor size roughly constant. Fully discriminating the form of PCD for each cell type and domain was challenging however, since both multiplexed imaging and spatial transcriptomics represent attempts to classify cells that are actively degrading their protein and mRNA components. We therefore present an aggregate analysis of the PCD in tumor cells, but provide the full Xenium dataset for follow up analysis by others.

### Non-proliferative and quiescent cancer cell states

To explore the role of quiescent cancer cells (QCCs) in MEL101, we quantified tumor cells that were negative for KI67 and positive for either or both of p21 (CDKN1A) and p27 (CDKN1B). By this definition, we found that QCCs represented 12% of all SOX10^+^ tumor cells, with the proportion varying from one tumor domain to the next. This is consistent with the presence of an MPI = –1 state in approximately 20% of SOX10^+^ tumor cells (**Supplementary Fig. 7a**; **Supplementary Fig. 4e**). We also performed gene signature analysis on both GeoMx and scRNA-seq data using quiescence gene signature^26^ and found that the magnitude of this signature was inversely correlated with that of proliferation signatures (G1/S and G2/M) (**Fig. 6a, b**; **Supplementary Fig. 7b, c**). The strength of quiescence signatures varied within MEL101 domains but was overall comparable to signature scores observed in progressing tumors MEL107–109 (**Supplementary Fig. 7d**). Thus, both assessments of cell cycle marker levels (measured by CyCIF) or gene signatures (inferred from GeoMx data) demonstrate that QCCs are present in MEL101 but are not a dominant tumor feature.

**Figure 6.**
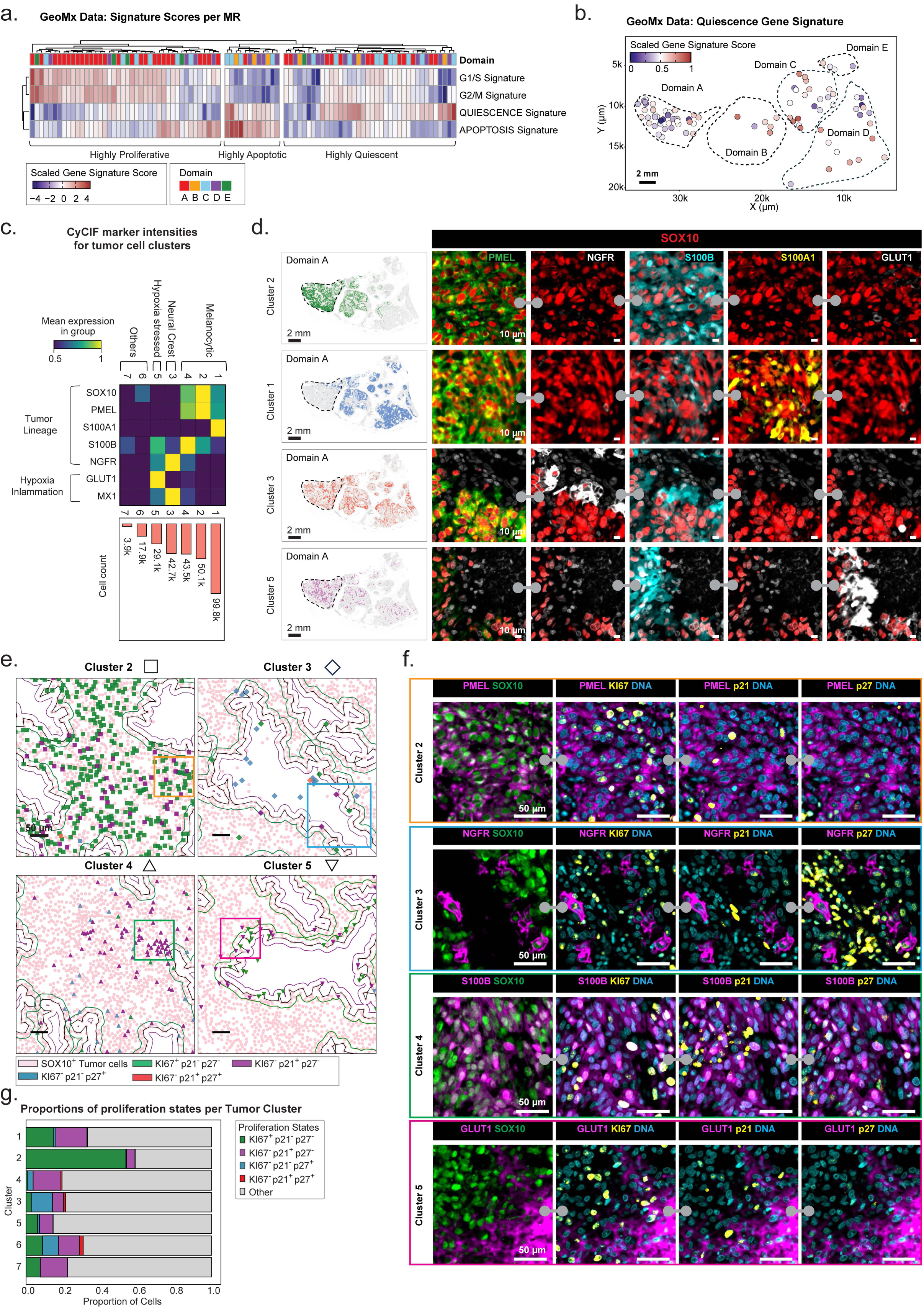
Presence of non-proliferative and quiescent cancer cell states in MEL101. **a.** Heatmap showing gene signature scores of gene signatures associated with proliferation, programmed cell death, and quiescence across all tumor MRs. Hierarchical clustering identified three dominant tumor states: highly proliferative, highly apoptotic, and highly quiescent. Columns represent tumor MRs annotated by Domains A–E (see color code below). **b.** Scatter plot showing spatially resolved QUIESCENCE signature scores across GeoMx tumor MRs. Each data point represents a tumor MR, and color indicates the scaled gene signature score. Scale bar, 2mm. **c.** Heatmap depicting the mean scaled expression of selected CyCIF markers for SOX10^+^ tumor clusters defined by Leiden clustering. The bar plot shows the absolute cell count for each cluster. **d.** Distributions of tumor cell states based on the clusters shown in panel *c*. *Left*: scatter plots depicting the spatial location of different tumor clusters, colored by cluster label, with background cells shown in grey. The black dashed region indicates Domain A. *Right*: selected channels from a 64-plex CyCIF image of regions of Domain A: SOX10 (red), PMEL (green), NGFR (white), S100B (cyan), S100A1 (yellow), and GLUT1 (white). Scale bars, 10 µm. **e.** Scatter plots of selected regions of Domain A showing the proliferation status of tumor cells in clusters 2–5. Each datapoint represents a tumor cell, shapes denote the clusters and color indicates the proliferation status as determined by p21 and p27 staining. The green, black and purple contour lines denote tumor margin. The regions outlined with colored rectangles are shown as CyCIF images in panel *f*. **f.** Selected channels from a 64-plex CyCIF images stained for SOX10 (green), PMEL (magenta), NGFR (magenta), S100B (magenta), GLUT1 (magenta), DNA (cyan), KI67 (yellow), p21 (yellow), and p27 (yellow), Scale bars, 50 µm. **g.** Stacked bar plot showing the proportions of KI67^+^p21^−^p27^−^ (green), KI67^−^p21^+^p27^−^ (purple), KI67^−^p21^−^p27^+^ (blue), KI67^−^p21^+^p27^+^ (red), and Other (grey) cells within each tumor cluster.

To determine if specific tumor lineages were enriched in QCCs, we performed unsupervised clustering of CyCIF markers (for all SOX10 and/or PMEL-positive cells, **Fig. 6c; Supplementary Fig. 7e**). This resulted in seven tumor cell clusters; cells in these clusters had differing prevalences across MEL101 tumor domains (**Supplementary Fig. 8b**). Clusters 1, 2, and 4 represented melanocytic states, characterized by the highest levels of SOX10 and PMEL expression (**Fig. 6c; Supplementary Fig. 7e**) but with differences in the expression of S100 proteins: clusters 2 and 4 were S100B-high and predominantly located in tumor Domains A and B, while cluster 1 was S100A1-high and most prevalent in other domains (**Fig. 6d, e; Supplementary Fig. 7f**). The S100 family has 21 members that are expressed at varying levels across most solid tumors^27^; expression of S100B is elevated in nevus cells relative to malignant melanoma, and the opposite is true of S100A1^28^. The functional significance of these differences remains unclear, but differential expression of S100 proteins are an effective means of identifying spatially patterned differences in melanoma cell states^28^. Cells in tumor cluster 3 had elevated NGFR expression, consistent with dedifferentiation into a neural crest-like state (**Fig. 6c, d**); these cells were mainly found in peritumoral, immune cell-dense regions (**Fig. 6d**). Cells in cluster 5 had lower expression of SOX10 and PMEL but high expression of the GLUT1 glucose transporter, NQO1 (NAD(P)H quinone oxidoreductase 1), and COX4 (a mitochondrial cytochrome c oxidase); all three proteins are associated with a metabolically active state and in some cases metastasis and poor prognosis (**Fig. 6c, d**).^29,30^ These cells were primarily found adjacent to necrotic regions, which represents a hypoxic environment sometimes associated with increased cancer aggressiveness (**Fig. 6e, f**).^31^

These data demonstrate the presence of spatially distinct melanoma lineages and microenvironments across MEL101. However, the key point is that all lineages and local environments that we have been able to identify comprised a mix of dividing and arrested cells with no evidence that quiescence was restricted to specific tumor states or spatial patterns (**Fig. 6f, g**). In particular, p27 positive and MPI = –1 cells were not obviously restricted to perinecrotic, perivascular, or other discernable niches in the tumor compartment. Thus, the prevalence and spatial distribution of p27-high and MPI = –1 QCC is insufficient to explain the maintenance of a tumor dormancy.

### Proliferating tumor cells and immune surveillance in a stable lesion involving scar tissue

ORION images of biopsies from the five additional specimens in the Buchbinder et al.^13^ cohort for which tissue was still available (MEL102 to MEL106; **Fig. 1a**) revealed the presence of SOX10^+^ tumor cells in specimen MEL105 but no other specimens; MEL106 contained only fibrotic tissue, and the other biopsy specimens contained fibrotic tissue with infiltrating Treg and CD8^+^ T cells, CD163^+^ macrophages, CD20^+^ B cells, and CD11C^+^ myeloid cells, (**Supplementary Fig. 8a**). MEL105 was scored previously as a complete pathological response based on inspection of H&E images^13^, but we identified 406 SOX10^+^ tumor cells out of a total of 40,670 segmented cells in the whole-slide image of the biopsy specimen (**Fig. 7a, b**). Remarkably, half of these tumor cells were KI67^+^, representing a higher proliferation index than observed in MEL101 or progressing tumors (**Fig. 7c**). Viable tumor cells were present in nests surrounded by diverse types of T and B cells, suggestive of active immune editing (**Fig. 7d, e**). We conclude that MEL105 is likely to represent another case of tumor persistence via mass dormancy with ongoing tumor cell proliferation counterbalanced by immune editing.

**Figure 7.**
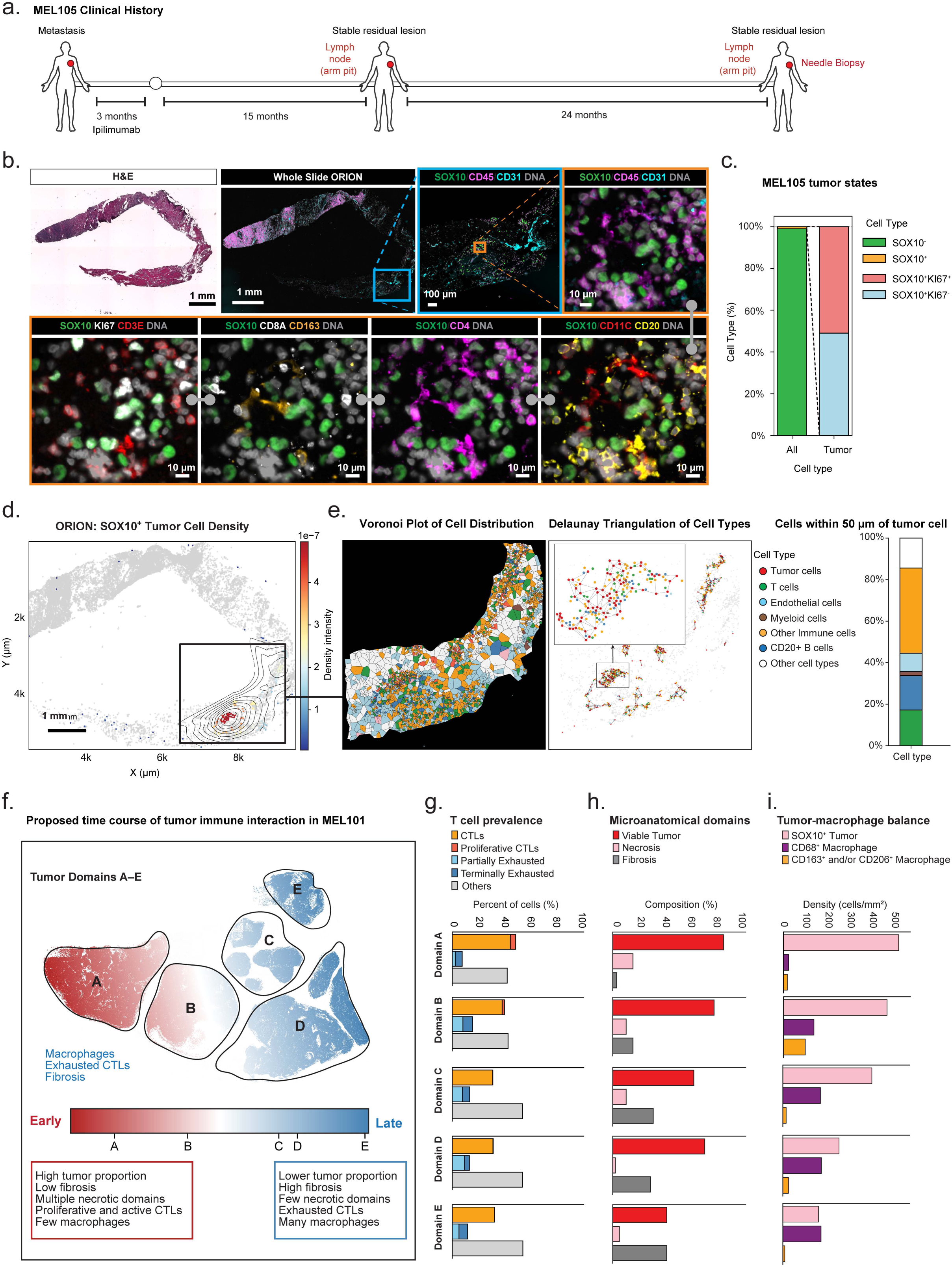
Proliferating tumor cells and immune surveillance in a stable lesion involving scar tissue. **a.** Clinical timeline for patient MEL105, whose tumor was biopsied after 24 months of stable disease. **b**. H&E-stained and ORION images of the needle biopsy specimen from MEL105. The grey barbell icon denotes the same ORION image with different channels shown. Selected channels from an 18-plex ORION images shown: SOX10 (green), CD45 (magenta), CD31 (cyan), DNA (grey), KI67 (white), CD3E (red), CD8A (white), CD163 (orange), CD4 (magenta), CD11C (red), CD20 (yellow). The regions outlined with blue and orange rectangles represent progressively magnified regions. Scale bars, 1 mm, 100 µm, and 10 µm. **c.** Stacked bar plot showing the percentage of tumor cells relative to all cells in the specimen (∼1%) and the proliferation status of these cells in specimen MEL105. **d.** KDE plot showing spatial distribution of SOX10^+^ tumor cells in specimen MEL105. Contour lines indicate regions of high local density, overlaid on all cells (grey). The color scale represents kernel density-estimated spatial intensity (range: 0–5 × 10^−7^, scaled unit). The region outlined with the black rectangle represents the magnified region in panel *e*. Scale bar, 1mm. **e.** *Left*: Voronoi plot showing the distribution of tumor, immune, and stromal cell types within the highlighted area in panel *d*. *Middle*: Delaunay triangulation plot of cell types surrounding SOX10^+^ tumor cells within a 50 µm radius. Black lines represent a Delaunay-based connectivity network between nearby SOX10^+^ tumor cells and surrounding immune cells. The region outlined with the black rectangle represents the magnified region. Each dot represents a single cell, colored by cell type. *Right*: stacked bar plot showing the percentage of cell types within a 50 µm radius of SOX10^+^ tumor cells; color codes match those in the Delaunay triangulation. **f.** Schematic representation of tumor-immune interactions across Domains A–E. **g.** Stacked bar plots showing the proportion of CD8⁺ T cell subsets out of all CD8^+^ T cells within Domains A–E. **h.** Bar plots showing the histopathologic composition of Domains A–E, as evaluated from H&E images. **i.** Bar plots showing the density (cells/mm^2^) of SOX10⁺ tumor cells, CD68⁺ macrophages and CD163⁺ and/or CD206⁺ macrophages across Domains A–E.

### Spatial variation in proliferation, death and dormancy

Variation in tumor and immune state across MEL101 is conceptualized in **Fig. 7f** as representing different phases of tumor-immune interaction. Domain A had the highest tumor and proliferating tumor cell density, in conjunction with the highest proportion of CTLs and proliferative CTLs, and highest level of necrosis (**Fig. 7g–i**). Domains B and C also had infiltrating CTLs, but also increasing proportions of macrophages, consistent with later stages of immune-mediated clearance of dead cells. Domains D and E were enriched in fibrosis, the terminal stage of tumor cell clearance. Thus, tumor mass dormancy in a single clinically stable tumor lesion is consistent with qualitatively distinct tumor states and phases of tumor-immune interaction.

## DISCUSSION

In this paper we use multimodal spatial profiling and conventional histology to study metastatic melanomas that exhibited persistent residual (stable) radiographic disease following anti-CTLA4 and/or anti-PD1 directed immunotherapy. We found that two out of six lesions contained actively dividing tumor cells but also infiltrating CTLs, consistent with active T-cell mediated tumor surveillance. One stable tumor (MEL101) was sufficiently large that it provided a unique opportunity to study tumor mass dormancy in detail in a human patient rather than a mouse model.^32^ Deep characterization of this tumor identified that the proportion of p27^+^ and/or p21^+^ QCCs in these tumors was similar to or lower than the proportion of proliferating cells. This demonstrates that the persistent tumors we examined are not primarily comprised of quiescent (non-dividing) cancer cells but have instead achieved a state of tumor mass dormancy involving ongoing cancer cell proliferation and balanced cell death. We found no evidence of tumor cells in the other four specimens, but biopsies contained viable immune cells, and it is likely that these contributed to the FDG signal detected on PET.

The observation that cells in MEL101 are as likely to express proliferation markers as tumors progressing on ICI therapy is striking. At a macroscopic level, the presence of necrosis, particularly in tumor Domain A, is *prima facie* evidence of ongoing cell death, most likely of both tumor and immune cells. Necrosis is often considered to be a negative prognostic feature^24,33^, but in MEL101 it is clearly compatible with effective immune-mediated tumor control. We were able to detect both individual dying tumor cells and fields of extensive death by CyCIF and to infer the activation of four different PCD pathways by GeoMx, scRNA-seq and Xenium data. Clearly distinguishing the mechanism of death has proven challenging in the crowded environment of an infiltrated tumor, likely because we are capturing cells in the middle of a process that results in degradation of the transcriptome and proteome. Specifically, our data suggest the involvement of cell-intrinsic PCD, potentially due to cell stress and metabolic deficiencies, but this is difficult to conclusively prove in the background of other PCD mechanisms and death of immune cells themselves. The high level of CTLs in MEL101, and the proximity of immune and tumor cells in peri-necrotic domains (given that necrosis is produced by cell death) argues that much of the tumor cell death we observe is immune-mediated. This suggests persistent anti-tumor immunity after ICI activation as a mechanism of durable tumor control and progression-free survival in these patients.

Recent research on persistent tumors in model organisms has focused on tumor cell quiescence and the factors that promote the QCC state. We detected cells with the properties of QCCs throughout MEL101, but these were not enriched in any previously described tumor states (e.g., dedifferentiated, neural crest-like, or melanocytic). Moreover, a p21 and/or p27 high QCC state was not the dominant tumor cell state, and it was not patterned spatially. This is suggestive of exit and entrance of cells from proliferation - a normal feature of cell division - rather than wide-spread entry into a dormant, quiescent, or senescent state. Our results therefore support earlier theories of tumor mass dormancy involving a dynamic balance between tumor cell division and cell death^4^.

Understanding the properties of dormant tumors and the sites they occur is of considerable clinical and biological interest since dormancy may be widespread. A long-term study of thin primary melanomas (T1) showed that 15–29% of patients had died from melanoma within 20 years of the original diagnosis, with the majority of deaths occurring >5 years after diagnosis^34^. Prior clinical evidence that the immune system plays a role in tumor dormancy is based primarily on transplantation cases in which immunosuppressed recipient patients developed melanoma after receiving organs from an immunocompetent donor who had a history of melanoma that was completely resected without evidence of existing disease.^35,36^ Other evidence of tumor-immune interactions leading to tumor dormancy have come primarily from pre-clinical models^6,37^. In these models, the immune system is observed to have a cytostatic effect on the tumor, resulting in a decrease in markers of tumor proliferation in the dormant vs progressive tumors. In contrast, our data reveal tumor proliferation in persistent stable lesions at the same or higher level than melanomas from patients with progressing lymph node tumors.

The dynamics of tumor mass dormancy within a single patient’s tumor are strikingly diverse and more varied than what has been observed in preclinical models. Within a single tumor, different regions display variable degrees of proliferating tumor cell density, immune cell activation, and cell death, all within a setting of thick fibrosis with no evidence of residual normal lymph node architecture (**Fig. 7f**). This suggests dynamism in the tumor-immune interaction, with regions of increased active tumor proliferation and anti-tumor immune response, co-existing with other relatively quieter regions. Fibrosis likely represents scars of past tumor-immune battles whereas necrosis is the more immediate sequela of tumor and immune cell death. Across MEL101, necrosis is most prominent where immune cell killing is greatest (in Domain A) and fibrosis where CTLs are least abundant, and macrophages enriched. Determining whether this reflects different stages of an anti-tumor immune response (as hypothesized in **Fig. 7f**) or differences in tumor clones is an important direction for future inquiry.

From a clinical perspective there is robust data that ICIs can provide long-term disease control in a substantial subset of advanced melanoma patients,^10,38^ but late progression (>5 year) in ICI-treated advanced melanomas has been reported to occur in 10–50% of responders depending on the setting and criteria.^39,40^ Thus, identification and management of patients at risk of progression is necessary to further improve outcomes. Our data suggest that a proportion of patients with persistent but stable residual lesions post-ICI may harbor viable residual tumor in those lesions. They also indicate that high FDG-avidity can be present in lesions with and without viable tumor cells, while low FDG-avidity can occur in lesions containing viable (and proliferative) tumor nests. These findings suggest that FDG-avidity is neither specific (e.g., due to fludeoxyglucose-18 uptake by proliferating non-malignant cells) nor sensitive (e.g., in small lesions) for identifying patients who would benefit from biopsy. Consistent with this observation, studies in the neoadjuvant setting have demonstrated that pathologic complete response associated with improved long-term outcomes and that radiographic responses do not always accurately reflect the underlying pathological response. The presence of reservoirs of viable and actively dividing tumor cells in persistent lesions is likely to have important implications for clinical monitoring and patient management. Residual viable disease would necessitate additional therapy, while residual lesions containing only scar tissue would not. However, it is noteworthy that patients in our cohort with detected viable tumor cells did not progress in eight years of subsequent follow-up, including the patient whose residual lymph node lesion (MEL105) contained viable melanoma cells and was only biopsied rather than completely resected. Importantly, our study suggests that tumor progression in this setting may involve immune escape rather than exit from a quiescent tumor cell state, and effective treatment strategies may thereby differ.

### Limitations

This study is necessarily limited by a relatively small number of patients, largely due to the challenge of identifying individuals who have post-ICI persistent lesions amenable for safe biopsy and enrolling them in a research study. While there are other potential sources of post-ICI samples including patients treated with neoadjuvant ICI therapy, the interval between treatment and surgery is too short in this setting to determine whether a tumor is clinically “dormant” (i.e. persistent and stable). Archived tissue from patients with melanoma treated with immunotherapy usually comes from symptomatic progressing lesions or regions of isolated progression which represents a different tumor state.

Nonetheless, we clearly identify a distinct subset of persistent residual tumor lesions with viable tumor cell proliferation and death in the post-ICI setting. Despite the small number of samples, our data suggests that a surprisingly substantial proportion of lesions in this clinical setting may have this phenotype. Broadly, our study supports the further study of tumor mass dormancy, as opposed to tumor cell quiescence, as a driver of tumor persistence in clinical cancer.

## Supporting information

Supplementary Table 1 COMBI-d cohort clinical and genomic data

Supplementary Table 2 Recurrent genomic alterations for BRAF-mutant melanoma

Supplementary Table 3 Gain of function alterations in comparison between DR and RP

Supplementary Table 4 Loss of function alterations in comparison between DR and RP

## ACKNOWLEDGEMENTS

We thank Jia-Ren Lin, Ajit Nirmal, Yu-an Chen, Daniel Lu, Michael Manos, Guihong Wan, Boshen Yan, Jihye Park, Amanda Garza, Kun Huang, Alyce Chen, Yilin Xu, Tyler Aprati, Rong Huang, Amy Huang, and Crystal Chiu for assistance and advice.

## AUTHOR CONTRIBUTION STATEMENTS

Y.S., Z.M., E.B., D.L., and P.K.S. conceived and designed the study. D.L., E.V.A., and P.K.S. supervised the project and acquired the funding. E.B. and P.O. supervised and provided clinical information for all the patients. S.S. and C.L. collected and harvested patient tissue blocks for the analysis. Y.S., Z.M., T.V., S.P., R.P., and B.K. collected and interpreted the molecular and multiplexed imaging data, with assistance from C.L. S.P. and Y.H. assisted with single-cell RNAseq analysis. Y.S., Z.M., E.B., D.L., and P.K.S. wrote the manuscript with input from all authors.

## DATA AND SOFTWARE AVAILABILITY

GeoMx gene expression data will be made available via the Gene Expression Omnibus (GEO; RRID: SCR_005012). All image data supporting the findings of this study are available via an index page on GitHub (https://github.com/labsyspharm/2025_Shi_Exceptional_Responder_Manuscript), which has been archived on Zenodo (http://doi.org/10.5281/zenodo.17807597). Code used for multimodal spatial analysis is available on GitHub.

## METHODS

### Clinical Samples

Patient samples from a recent publication (Buchbinder et al)^13^ or from patients with a medical history of melanoma progression after ICI treatment (Supplementary Table 1) were retrieved from the archives of the Department of Pathology at Brigham and Women’s Hospital under BWH Institutional Review Board approval (Protocol 05-042), under a waiver of consent. Following de-identification, spatial profiling of these specimens was approved under HMS Institutional Review Board Protocol IRB21-0656. Fresh 5 µm sections were cut from each tumor block. The first section of each block was H&E stained for histopathological review and annotation of the sample. The remaining sections were characterized using immunohistochemistry (IHC), cyclic immunofluorescence (CyCIF, ORION) microscopy and microregional whole-transcriptome profiling (Nanostring GeoMx).

Treatment history and clinical metadata are shown in **Supplementary Table 1**. In brief, patient **MEL101** was diagnosed with metastatic melanoma in 2013, with lesions in the brain, left axilla, supraclavicular lymph nodes, and adrenal glands. The patient received four cycles of ipilimumab (anti-CTLA-4 blockade), resulting in a response and resolution of all lesions except for a persistent residual lymph node metastasis in the left axilla. At the time of this analysis (17 years post-initiation of ipilimumab), the patient remains free of recurrent disease. The clinical courses for patients **MEL102–MEL106** are described in the previous publication.^13^ Patient **MEL107** was diagnosed with metastatic melanoma involving the lungs and initially treated with pembrolizumab without clinical response. Subsequent administration of two doses of ipilimumab resulted in a complete response. Eighteen months later, a recurrent right axillary lymph node metastasis was resected and included in this study. Patient **MEL108** received initial treatment consisting of surgical resection and a tumor vaccine. One year later, the patient experienced recurrence and was treated with ipilimumab and nivolumab, yielding a mixed response. A non-responding left axillary lymph node was resected and included in this study. The patient ultimately died of melanoma five years after the initial diagnosis. Patient **MEL109** presented with nodal metastatic melanoma and underwent cervical lymph node resection followed by ipilimumab. The patient progressed 4 months later and was started on pembrolizumab, which was subsequently held due to pneumonitis. Ten months after initiating pembrolizumab, she again showed disease progression and was re-started on pembrolizumab, achieving a complete response and completing two years of therapy. Ten months after stopping pembrolizumab, she developed progression in an inguinal lymph node, and the biopsy of that node was used for this study.

### Tissue imaging (H&E, IHC, CyCIF)

H&E and IHC staining of FFPE sections was performed by the Brigham and Women’s Hospital (BWH) Pathology Core and digitized using an Olympus VS-120 automated microscope using a 20x/0.75 NA objective at the Neurobiology Imaging Core at Harvard Medical School. IHC for cleaved caspase-3 used rabbit monoclonal antibody (clone A5E1, 1:500 dilution, Cell Signaling Technologies) followed by HRP-based chromogenic detection and hematoxylin counterstain. CyCIF was performed for specimens MEL101 and MEL107–MEL109 as previously described^14^ and at protocols.io (dx.doi.org/10.17504/protocols.io.j8nlkoqbdv5r/v1). In brief, the BOND RX automated slide stainer was used to bake slides at 60 °C for 30 minutes, dewax using Bond Dewax solution at 72 °C and perform using Epitope Retrieval 1 (Leica) solution at 100 °C for 20 minutes. After 4 rounds of photochemical bleaching to reduce tissue autofluorescence, slides underwent multiple cycles of antibody incubation, imaging, and fluorophore inactivation. Tissues were incubated overnight in the dark at 4 °C in antibody solution containing Hoechst. Coverslips were mounted with 70% glycerol 1x PBS prior to imaging with a CyteFinder slide-scanning fluorescence microscope (RareCyte Inc.) with a 20x/0.75 NA objective. Slides were soaked in 42 °C PBS to facilitate coverslip removal, and then fluorophores were inactivated by photochemical bleaching in a tray containing 4.5% H_2_O_2_ in PBS supplemented with 24 mM NaOH placed on a LED light source for one hour at room temperature. The list of all the antibodies and their dilutions for each experiment is presented in **Supplemental Table 3**.

### Tissue imaging (Orion)

Tissue imaging with RareCyte ORION platform (RRID:SCR_026855) was performed for specimens MEL102–MEL106 with following protocol previously described by Lin et al.^15^ In brief, tissue sections were deparaffinized and subjected to heat-induced epitope retrieval. Slides were incubated at 95 °C for 5 min and then at 105 °C for 5 min in EZ-AR2 Elegance buffer (BioGenex HX032). Autofluorescence was reduced by treatment with a peroxide-based quenching solution (4.5% H₂O₂, 25 mM NaOH in PBS) under illumination, followed by washes in surfactant-containing buffer. Slides were then incubated in ORION Signal Enhancer for 15 min, washed, and stained overnight at 4 °C with the multiplex antibody panel. After primary staining, sections were washed and incubated for 30 min with anti-DIG antibody and streptavidin to detect DIG- and biotin-labeled probes. Slides were washed and coverslipped with ORION mounting medium (RareCyte WA). For sequential H&E staining, ORION-stained sections were decoverslipped in PBS for overnight, followed by the standard H&E protocol.

### CyCIF and ORION image pre-processing and quality control

Assembly of raw CyCIF and ORION imaging data into a high-dimensional images, as well as nuclear segmentation and single-cell feature extraction, were performed using the open-source MCMICRO^41^ processing pipeline (RRID:SCR_022832), an open-source multiple-choice microscopy pipeline (version:38182748aa0ec021f684ce47248c57340d2f4cc7; full codes available at https://github.com/labsyspharm/mcmicro). The specific parameters used were optimized after iterative inspection of results, specifically focusing on performance of the segmentation module to ensure accurate identification of single cells (params.yml files available at https://github.com/labsyspharm/2025_Shi_Exceptional_Responder_Manuscript). The quality of the segmentation was assessed, and the segmentation parameters were iteratively modified to improve the accuracy of the segmentation masks. After generating the segmentation masks, the mean fluorescence intensities of each marker for each cell were computed, resulting in a single-cell data table for each acquired whole-slide CyCIF and ORION image. The X/Y coordinates of annotated histologic regions on the whole slide image were used to extract the single-cell data of cells that lie within the annotated regions. Multiple approaches were also taken to ensure the quality of the single-cell data. At the image level, the cross-cycle image registration and tissue integrity were reviewed; regions that were poorly registered or contained severely deformed tissues and artifacts were identified, and cells inside those regions were excluded. Antibody staining patterns that gave low confidence by visual evaluation were also excluded from the analyses.

### CyCIF and ORION single-cell gating-based phenotyping

Cells from CyCIF and ORION data were phenotyped using a gating-based classification approach as previously described.^42^ In brief, gates for each marker were defined using an open-source visualization tool (https://github.com/labsyspharm/gater). These gates were then used to rescale single-cell expression values between 0 and 1, with values above 0.5 indicating marker positivity. The normalized data were subsequently used for cell-type calling and unsupervised clustering. Phenotype labels were assigned using the SCIMAP^43^ Python package (RRID:SCR_024751) based on a hierarchical classification of marker expression patterns (**Supplementary Fig. 2a**). Assigned cell types were validated by overlaying phenotype labels onto the corresponding images.

### Definition of microanatomical regions

Microanatomical regions (tumor, peritumoral, necrotic, and perivascular) were defined using the SpatialCells package (https://github.com/SemenovLab/SpatialCells).^44^ Single-cell centroid coordinates of SOX10^+^ tumor cells were used to delineate tumor boundaries by applying density-based spatial clustering (DBSCAN) followed by polygon-based boundary reconstruction to defined contiguous tumor regions. Regions immediately adjacent to tumor boundaries were classified as peritumoral regions. Necrotic regions were subclassified within peritumoral region based on the presence of DNA debris staining and sparse cell counts. Perivascular regions were defined as areas proximal to CD31^+^ endothelial cells. After region assignment, cell-type proportion within each microanatomical regions (shown in **Supplementary Fig. 3d**) were calculated by normalizing the number of cells of each cell type to the total number of that cell type across Domains A–E.

### Clustering and UMAP (CyCIF data)

To define tumor cell states from CyCIF imaging data, SOX10^+^ tumor cells were filtered from the full dataset MEL101A Level 1, Slide 2 and analyzed using the SCIMAP^43^ package in Python. Expression intensity of all the protein markers was normalized based on gating thresholds and used to construct a k-nearest neighbor graph. Dimensionality reduction was performed using UMAP, and unsupervised clustering was conducted using the Leiden clustering with a resolution of 0.3 to identify distinct tumor clusters.

### Multivariate Proliferation Index (MPI) calculation

The Multivariate Proliferation Index (MPI) was based on the normalized measurement of five markers: three proliferation markers (KI67, CCNA2, CCNB1) and two cell-cycle arrest markers (p21, p27). The method avoids relying on single markers while separating cells expressing high level arrest markers (even if proliferation markers are expressed). The detailed methodology can be found in Gaglia, G. et al.^18^ In the original study, cells with an MPI score of −1 were classified as being a cell cycle-arrested state. However, subsequent study has suggested that p27 is a more specific marker of cell-cycle arrest than p21.^17^ Therefore, while MPI scores were calculated using the same framework as described by Gaglia, G. et al,^18^ we applied a more stringent classification scheme for proliferation status. Specifically, cells with MPI = 1 were classified as proliferative, cells with MPI = −1 were classified as non-proliferative, and all remaining cells were assigned to the “Others” category (MPI = 0).

### CyCIF kernel density estimation and contour mapping

For each phenotypic population, spatial density maps were generated using kernel density estimation (KDE) on single-cell centroid coordinates. All cells within each field were plotted as low-opacity background points to provide spatial context. For each phenotype, cells were selected and, when the number of cells was at least 50, a Gaussian KDE was applied to the centroid coordinates of marker positive cells with intensity clipped at the 98^th^ percentile and regions below the 10^th^ percentile masked for contrast enhancement. Density values were evaluated at cell positions and interpolated onto a regular two-dimensional grid to generate continuous density surfaces. Upper density values were clipped, and lower-density regions were masked and colored according to KDE-derived density intensity.

### CyCIF neighborhood spatial analysis

Cell centroid coordinates were converted to microns. A Delaunay triangulation was performed on the coordinates of the target cell population, and edges were adaptively filtered based on local nearest-neighbor distance to retain short-range connections. Cells located within 50 μm of any target cell were defined as neighbors. In **Fig. 2c** (perivascular region), CD8^+^ T cells and CD31^+^ endothelial cells were highlighted, and the filtered CD31^+^ connectivity network was visualized as line segments.

### CyCIF proximity analysis

CyCIF proximity analyses were performed using the *spatial_distance* function in the SCIMAP python package.^43^ For each single cell in the CyCIF data, the average distance to specific cell phenotypes or clusters was calculated in two-dimensional coordinate space, enabling quantitative assessment of spatial relationships among cell populations within tissue sections.

### Microregion transcriptomics (GeoMx^Ⓡ^) sample processing, data collection, and annotation

Digital spatial transcriptomic profiling was performed using the NanoString GeoMx^Ⓡ^ Human Whole Transcriptome Atlas (WTA) RNA probes, following previously published protocols.^45,46^ Briefly, freshly cut (< 2 weeks old) 5-µm thick FFPE melanoma sections were baked at 60 °C for 3 hours, dewaxed, and hybridized overnight with WTA probes. The following day, slides were incubated with fluorescence-conjugated antibodies targeting melanocytes (SOX10, 1:100, Abcam ab270151; MART1, 1:200, Abcam ab225500), and immune cells (CD45) before imaging and transcript collection on DSP (**Supplementary Fig. 2c**, **Supplementary Table S3**). These fluorescent marker signals were integrated with histopathology guides for MR selection. A total of 236 MRs were selected to present morphologically distinct zones, including the tumor edge, tumor center, peritumoral stroma, and tertiary lymphoid structure. Collected MRs were pooled for downstream library preparation and transcriptomic analysis.

### GeoMx analysis

GeoMx RNA sequencing data were processed using the Bioconductor GeoMxTools RNA-NGS workflow (https://www.bioconductor.org/packages/release/workflows/vignettes/GeoMxWorkflows/inst/doc/GeomxTools_RNA-NGS_Analysis.html). Raw count data were imported, quality controlled, and normalized according to the recommended pipeline, including probe-level filtering and sample-level quality control. Normalized expression matrices were used to perform t-SNE dimensionality reduction to visualize MR-level transcriptional heterogeneity.

### GeoMx gene signature analysis

Gene signature analysis was performed using DESeq2-normalized counts (obtained vis *counts(dds, normalized = TRUE)*) followed up log10 transformation. Single-sample gene set enrichment analysis (ssGSEA) scores were calculated using the GSVA R package (https://bioconductor.org/packages/release/bioc/GSVA/; RRID:SCR_021058), and pathway activity scores were used for downstream comparative analyses (e.g., **Fig. 3i**). Pathways included G1/S,^46^ G2/M,^46^ APOPTOSIS (Hallmark),^47^ NECROPTOSIS (GO:BP),^48^ PYROPTOSIS [Reactome],^49^ FERRORPTSIS (WP),^50^ and QUIESCENCE.^26^

### Single-cell RNA-seq analysis

Single-cell RNA-seq data was generated using 10x Genomics Chromium platform and processed with Cell Ranger^51^ (RRID:SCR_017344) for alignment, demultiplexing, and gene-cell count matrix generation. Downstream analyses were performed in Python using Scanpy^52^ (RRID:SCR_018139). Cells with low UMI counts, low gene detection, or high mitochondrial transcript content were filtered, and genes expressed in only few cells were removed. Data were normalized, log-transformed, and highly variable genes were selected. Dimensionality reduction was performed using PCA, followed by neighborhood graph construction, UMAP embedding, and Leiden clustering. Clusters were annotated into major cell types based on canonical marker gene expression.

### RNA velocity analysis

RNA velocity was computed from spliced and unspliced read counts generated using Velocyto^53^ (RRID:SCR_018167). Downstream analysis was performed with the scVelo package^54^ (RRID:SCR_018168). Moments were calculated on the k-nearest neighbor graph, velocities were estimated using the dynamic model, and a velocity graph was constructed. Velocity streamlines were projected onto the UMAP embedding and visualized over cell-type labels to infer transcriptional trajectories, with particular focus on state transitions within CD8^+^ T cells subsets.

### Single-cell RNA differential expression and gene set enrichment analysis

Differential gene expression of tumor cell clusters in **Fig. 3c** was performed using *rank_genes_groups* function from Scanpy.^52^ For tumor cluster 4, the top 100 upregulated genes ranked by log fold change were selected for downstream enrichment analysis. Functional enrichment was performed using gene ontology–based pathway analysis, and enrichment scores were calculated. Dot plots were generated to visualize significantly enriched biological processes, where dot size represents the percentage of genes in each gene set and color indicates –log10-transformed p-values.

### Xenium sample processing

FFPE tissue sections (5 μm) were cut from archival tissue blocks and processed using the Xenium Prime 5K platform (10x Genomics) according to the manufacturer’s standard protocol at the Molecular Imaging Core from Dana-Farber Cancer Institute. Following Xenium imaging and transcript detection, tissue sections were recovered and subject to CyCIF staining as described above.

### Xenium Analysis

Xenium Prime 5K data were processed in Scanpy. Gene expression matrices were imported from Xenium cell-feature matrices and integrated with cell-level metadata and spatial coordinates. The analysis was restricted to the MEL101 block 2 domain A annotated on histopathologist by selecting cells within manually annotated boundaries. Downstream analyses were performed in Python using Scanpy^52^ (RRID:SCR_018139). Cells with fewer than 20 total transcripts or transcripts counts above the 98^th^ percentile were excluded, and genes detected in fewer than 100 cells were removed. Data were normalized, log-transformed, and highly variable genes were selected. Dimensionality reduction was performed using PCA, followed by neighborhood graph construction, UMAP embedding, and Leiden clustering. Clusters were annotated into major cell types based on canonical marker gene expression. PCD programs were quantified at the single-cell level using module scoring (*scanpy.tl.score_genes*). Cells with program score at or above the 99^th^ percentile of the corresponding score distribution were classified as PCD-high and included in subsequent analyses.

### Statistical Tests

All statistical comparisons between groups were performed using the Mann-Whitney U rank test to assess differences in the distribution of values between groups, implemented via the *mannwhitneyu* function in the SciPy^55^ Python package (RRID: SCR_008058) or the stats and ggpubr R packages (RRID:SCR_021139).

