## Supplementary Figures for "Immunogenic tumor mass dormancy as a driver of persistent residual lesions in immunotherapy-treated melanoma"

### Supplementary Figure 1.

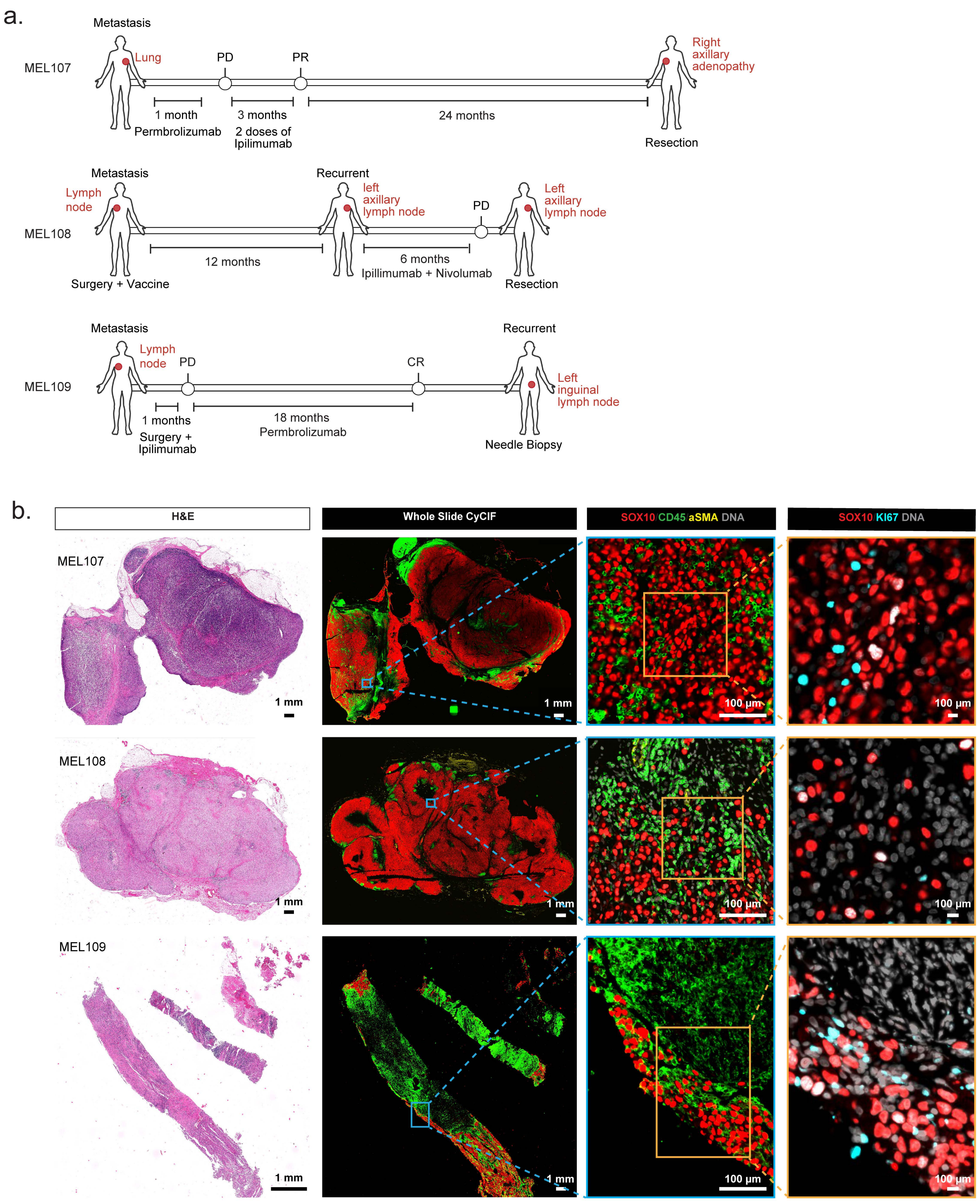

**Supplementary Figure 1. a.** Clinical timeline of patients MEL107–MEL109 (These schematics provide simplified representations of patients' treatment and progression histories; detailed clinical information is provided in METHODS). The red dots represent tumor locations. **b.** H&E-stained (left) and CyCIF (right) images of specimens MEL107–MEL109. CyCIF images are stained for SOX10 (red), CD45 (green), αSMA (yellow), and KI67 (cyan). Scale bars, 1 mm, 100 μm, and 10 μm.

Supplementary Figure 2.

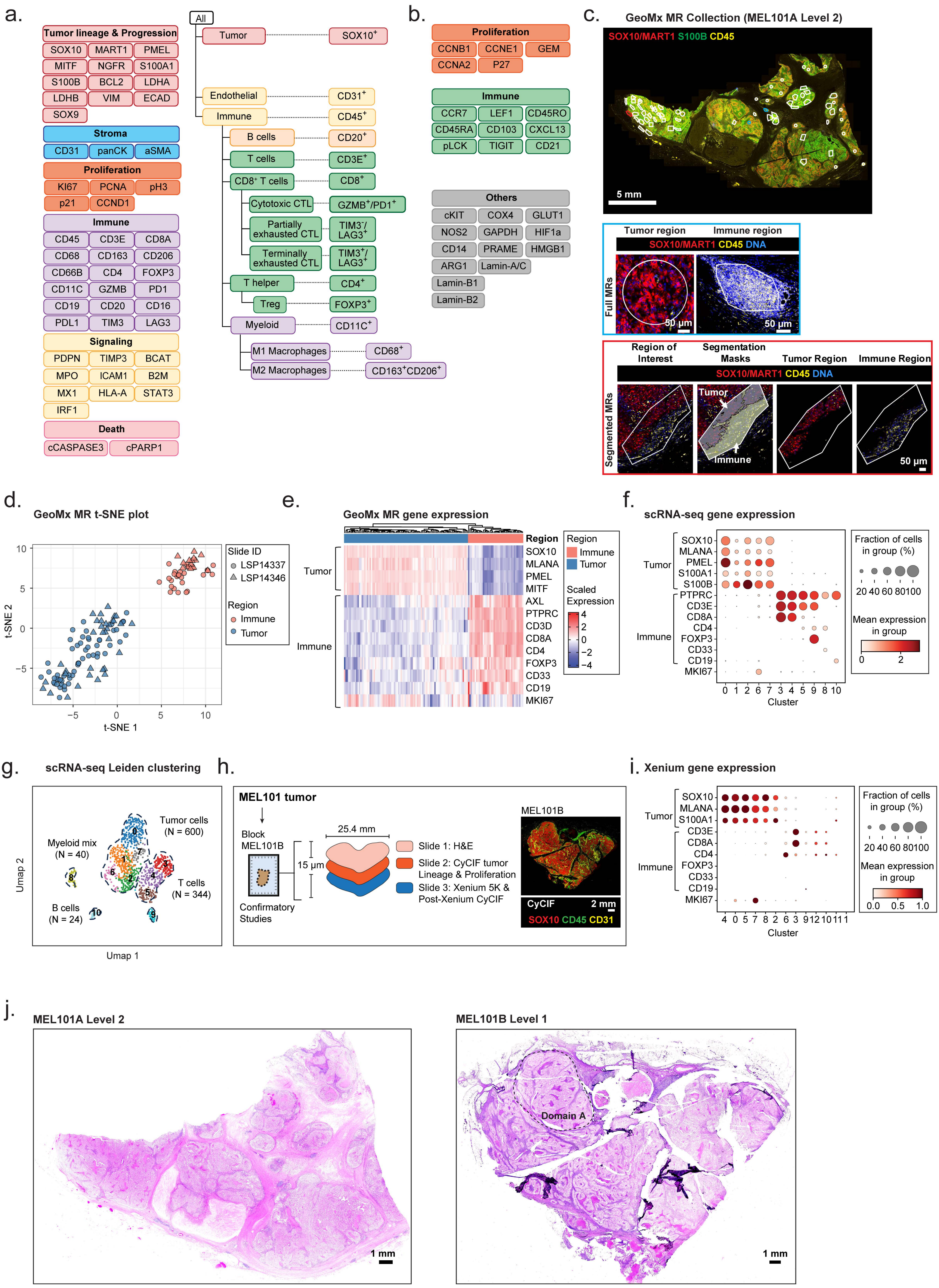

**Supplementary Figure 2. a.** Antibodies included in CyCIF panels listed in **Fig. 1c** (left). Flowchart of cell type calling strategy used for CyCIF data. **b.** Additional antibodies for cell cycle and T cell subtypes. **c.** CyCIF image of MEL101A Level 2 stained for SOX10/MART1 (red), S100B (green), and CD45 (yellow) (top). The blue and orange rectangles, illustrating full and segmented GeoMx MRs, are magnified below. Scale bars, 5 mm and 50 µm. **d.** t-SNE projection of GeoMx MR transcriptomes. MRs are annotated by region type and slide ID. The slide IDs represent two different batches of GeoMx collection from specimen MEL101. **e.** Heatmap of representative marker genes distinguishing tumor and immune MRs. Each column represents an individual MR. **f.** Dot plot of scRNA-seq Leiden clusters. The dot plot displays the expression of tumor- and immune-associated genes across clusters 0–10. Dot size reflects the percentage of cells expressing each gene, while color intensity represents the average normalized expression level. **g.** UMAP projection of scRNA-seq data showing major cell populations annotated based on Leiden clustering. **h.** Schematic of tumor specimens and multimodal experimental approach. FFPE block MEL101B was used in confirmatory studies to check overall tumor architecture, which generates multiple 5-micron sections for H&E, CyCIF and 10x Genomic Xenium 5K spatial transcriptomics. **i.** Dot plot of Xenium 5K Leiden clusters. The dot plot displays the expression of tumor- and immune-associated genes across clusters 0–10. Dot size reflects the percentage of cells expressing each gene, while color intensity represents the average normalized expression level. **j.** H&E-stained sections of specimens MEL101A Level 2 (left) and MEL101B Level 1 (right). The black dashed outline highlights Domain A in the MEL101B Level 1, identified based on histopathology. Scale bars, 1 mm.

Supplementary Figure 3.

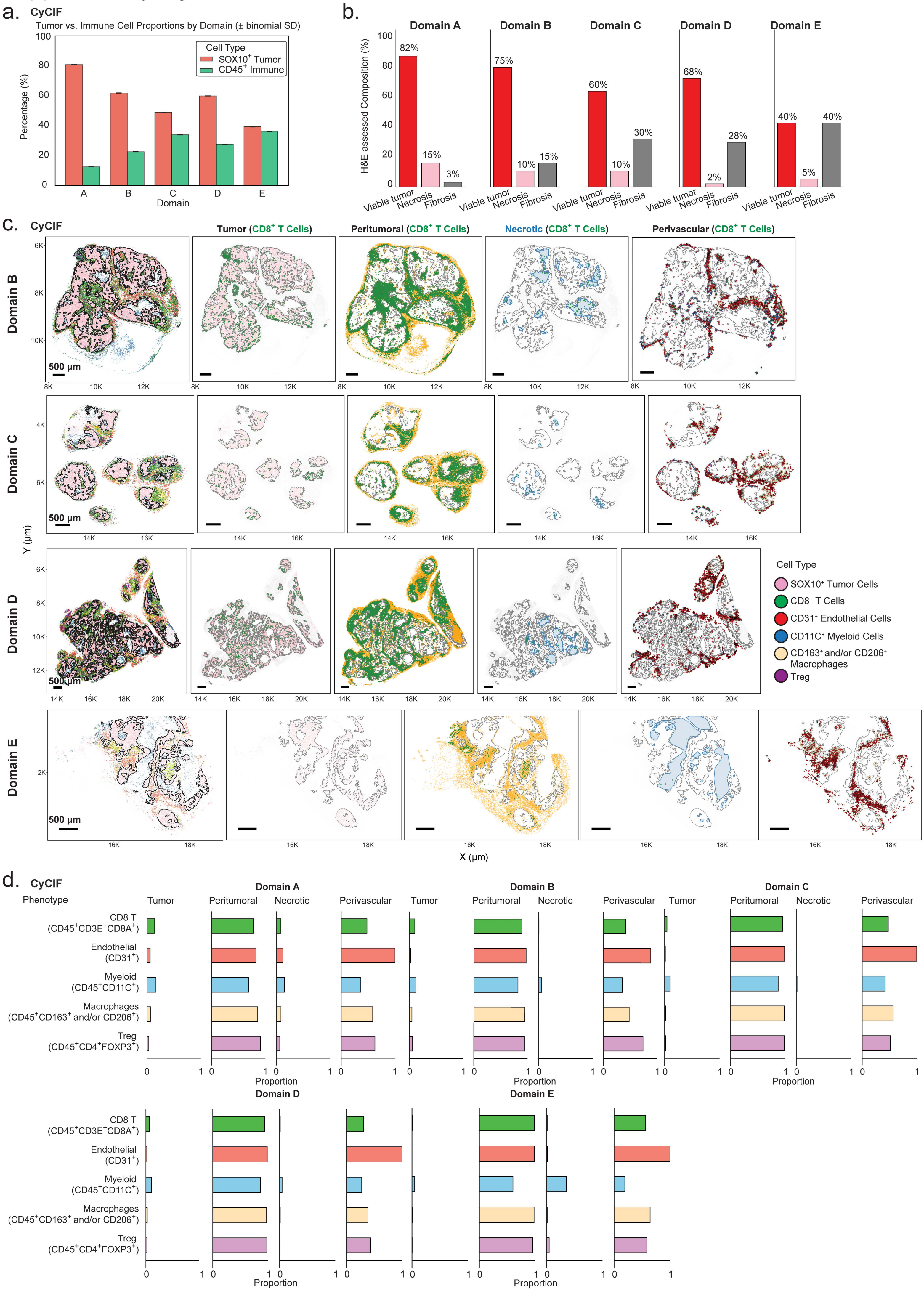

**Supplementary Figure 3. a.** Bar plot showing the percentage of $SOX10^+$ tumor cells and $CD45^+$ immune cells out of all cells within Domain A–E. Error bars represent binomial standard deviations. **b.** Bar plots showing the percentage of tumor mass, necrosis, and fibrosis assessed by H&E across all the domains. **c.** Scatter plot displaying the spatial distribution of selected cell types within Domains B–E (first column). The black outline denotes tumor boundary defined based on $SOX10^+$ tumor cells. The scatter plots in the remaining columns show the spatial distribution of $CD8^+$ T cells (green) within the microanatomical regions—Tumor (pink), Peritumoral (orange), Necrotic (blue) and Perivascular (red) regions—across Domains B–E. In the perivascular region, the black lines represent a Delaunay-based connectivity network between nearby $CD31^+$ cells (red dots) and $CD8^+$ T cells. Scale bars, 500 μm. **d.** Bar plots representing the distribution of various immune cell types within microanatomical regions across Domains A–E.

Supplementary Figure 4.

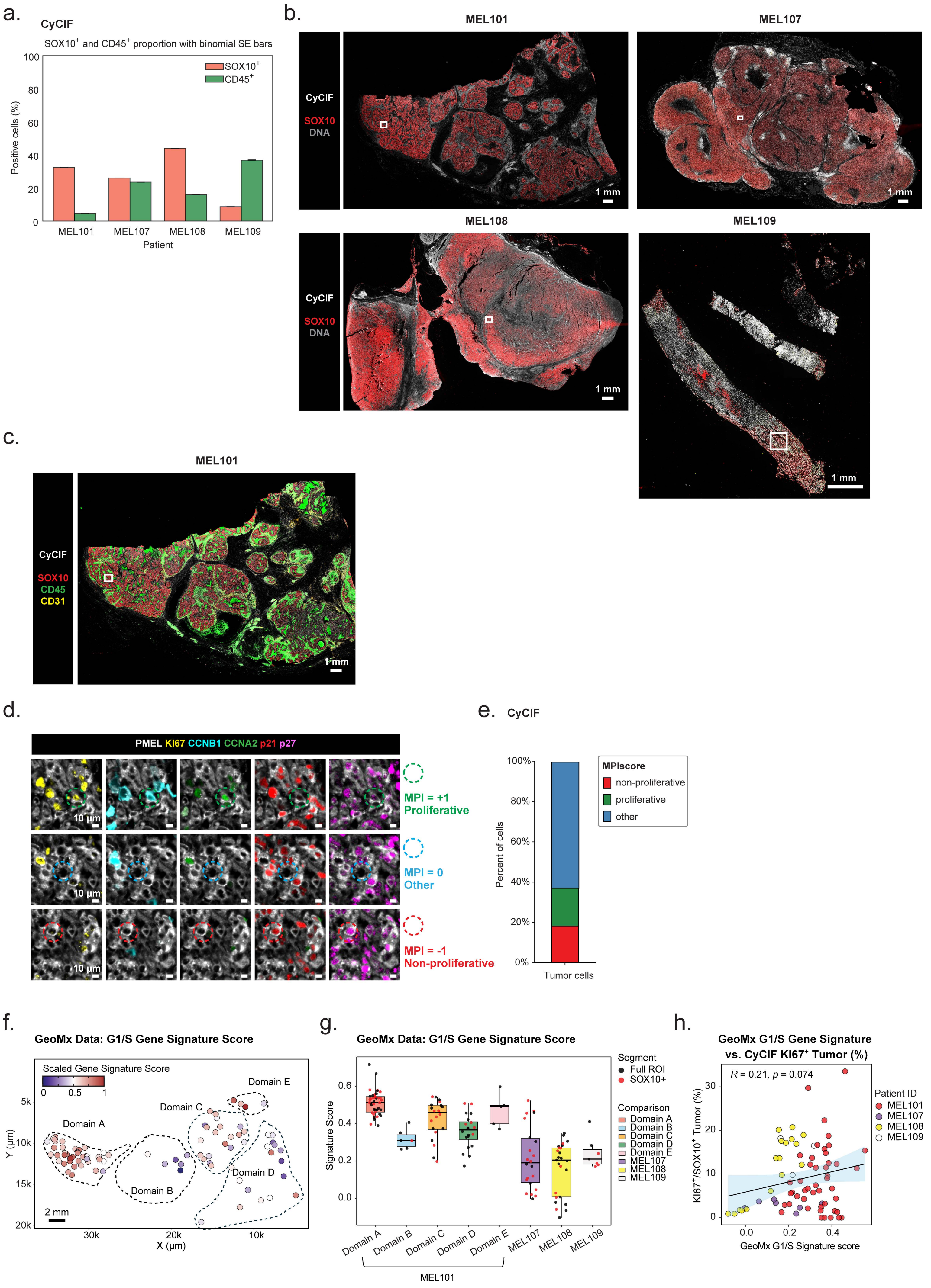

**Supplementary Figure 4. a.** Bar plot showing the proportion of $SOX10^+$ tumor and $CD45^+$ immune cells out of all cells within specimens MEL101 and MEL107–MEL109. **b.** CyCIF images of MEL101 and MEL107–MEL109 stained for SOX10 (red) and DNA (grey). The regions outlined with white rectangles are magnified regions shown in **Fig. 3a**. Scale bars, 1mm. **c.** CyCIF image of MEL101A Level 1 stained for SOX10 (red), CD45 (green), and CD31 (yellow). The region outlined with a white rectangle corresponds to the magnified region in **Fig. 3f**. Scale bar, 1 mm. **d.** CyCIF image of MEL101A Level 1 stained for KI67 (yellow), CCNB1 (cyan), CCNA2 (green), p21 (red), and p27 (magenta). Examples of proliferative (MPI = +1), non-proliferative (MPI = –1), and other (MPI = 0) tumor cells are highlighted with green, red, and cyan dashed circles, respectively. Scale bars, 10 µm. **e.** Stacked bar plot showing the proportion of proliferative (MPI = +1), non-proliferative (MPI = –1), and other (MPI = 0) tumor cells in MEL101. **f.** Scatter plot showing spatially resolved G1/S gene signature scores across GeoMx tumor MRs. Each data point represents a tumor MR, and color indicates the scaled gene signature score. Scale bar, 2mm. **g.** Boxplot comparing G1/S gene signature scores across tumor GeoMx MRs in Domains A–E and progressing tumors (MEL107–MEL109). Each dot represents a tumor MR (either $SOX10^+$ segmented or full tumor MRs). Boxes represent the first and third interquartile of the data; whiskers extend to show the rest of the distribution except for points identified as outliers. **h**. Scatter plot showing the Pearson correlation between the G1/S gene signature scores (x-axis, GeoMx) and the percentage of $SOX10^+KI67^+$ tumor cells among all tumor cells (y-axis, CyCIF). Each data point represents an individual full tumor MR colored by patient ID. The solid line indicates the linear regression fit, with a 95% confidence interval shown (shaded area). Significance was calculated using a two-sided Pearson correlation test.

Supplementary Figure 5.

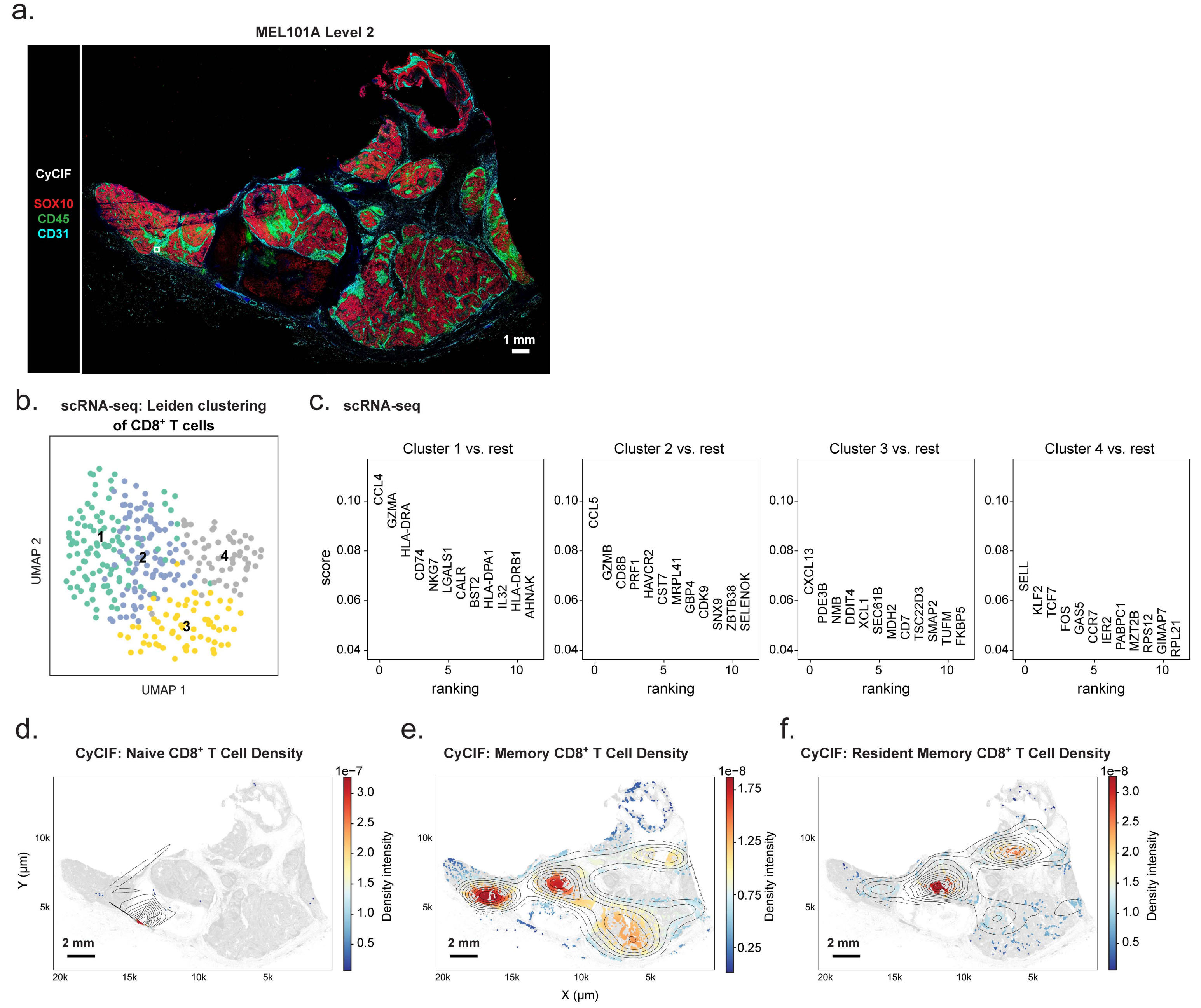

**Supplementary Figure 5. a.** CyCIF image of MEL101A Level 2 stained for SOX10 (red), CD45 (green), and CD31 (cyan). The region outlined with a white rectangle corresponds to the magnified region in **Fig. 4b**. Scale bar, 1mm. **b.** UMAP of $CD8^+$ T cells colored by $CD8^+$ T cell cluster ID (clusters 1–4), identified by Leiden clustering in scRNA-seq data. **c.** Top-ranked genes distinguishing the $CD8^+$ T cell clusters 1–4 from all other clusters (scRNA-seq data). The x-axis shows the rank order of the identified genes, and the y-axis indicates the corresponding gene expression scores. **d–f.** KDE plots showing the spatial distribution of specific $CD8^+$ T cell subtypes across MEL101 domains. Contour lines indicate regions of high local density, overlaid on all cells (grey). The color scale represents kernel density-estimated spatial intensity (range in scaled units as shown). **d.** Naïve $CD8^+$ T cells. **e.** Memory $CD8^+$ T cells. **f.** Resident Memory $CD8^+$ T cells. Scale bar, 2 mm.

Supplementary Figure 6.

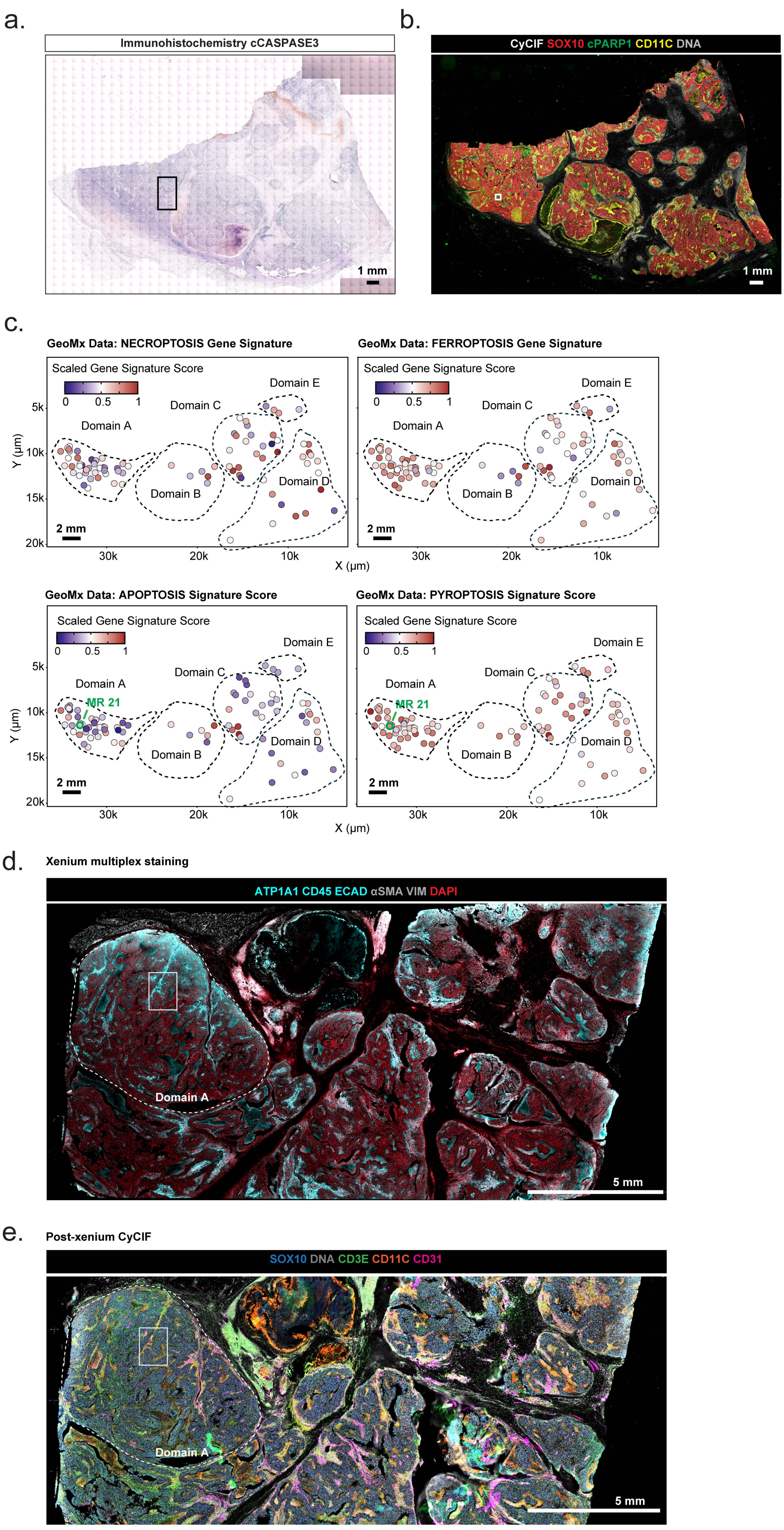

**Supplementary Figure 6. a.** cCASPASE3 immunohistochemistry image of specimen MEL101A Level 1. The region outlined with a black rectangle corresponds to the magnified region shown in **Fig. 5a**. Scale bar, 1mm. **b.** CyCIF image of MEL101A Level 1 stained for SOX10 (red), cPARP1 (green), CD11c (yellow), and DNA (grey). The region outlined with a white rectangle corresponds to the magnified region shown in **Fig. 5b**. Scale bar, 1mm. **c.** Scatter plot showing spatially resolved NECROPTOSIS, FERROPTOSIS, APOPTOSIS, and PYROPTOSIS signature scores across GeoMx tumor MRs. Each data point represents a tumor MR, and color indicates the scaled gene signature score. Scale bars, 2 mm. **d.** Representative Xenium imaging from MEL101B showing ATP1A1 (cyan), CD45 (cyan), ECAD (cyan), αSMA (grey), VIM (grey), and DAPI (red). The white dashed outline highlights Domain A in the MEL101B block, identified based on histopathology. The region outlined within a white rectangle corresponds to the magnified region shown in **Fig. 5g**. Scale bar, 5 mm. **e.** Post-Xenium CyCIF image of MEL101B stained for SOX10 (blue), DNA (grey), CD3E (green), CD11C (orange), and CD31 (red). The white dashed outline highlights Domain A in the MEL101B Level 1, identified based on histopathology. The region outlined within a white rectangle corresponding to the magnified region shown in **Fig.5f, g**. Scale bar, 5 mm.

Supplementary Figure 7.

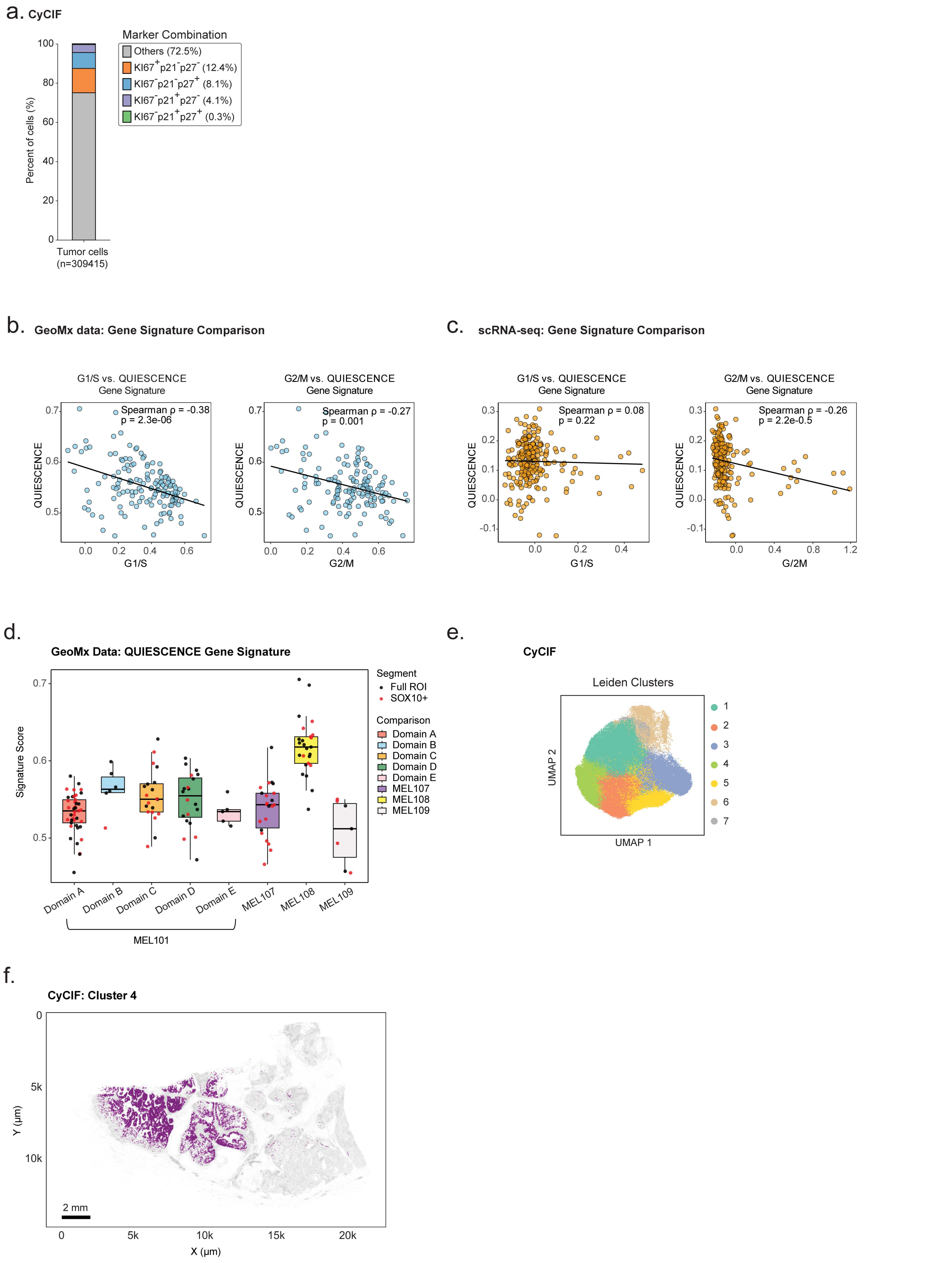

**Supplementary Figure 7. a.** Stacked bar plot of $KI67^{+}p21^{-}p27^{-}$, $KI67^{-}p21^{+}p27^{-}$, $KI67^{-}p21^{-}p27^{+}$, $KI67^{-}p21^{+}p27^{+}$, and Others proportion of tumor cells in specimen MEL101. **b.** Scatter plot showing the Pearson correlation between the G1/S (left) and G2/M (right) (x-axis) and QUIESCENCE gene signature scores (y-axis). Each data point represents an individual tumor MR. The solid line indicates the linear regression. Significance was calculated using a two-sided Spearman correlation test. **c.** Scatter plot showing the Pearson correlation between the G1/S (left) and G2/M (right) (x-axis) and QUIESCENCE gene signature scores (y-axis). Each data point represents a single cell from scRNA-seq data. The solid line indicates the linear regression. Significance was calculated using a two-sided Spearman correlation test. **d.** Boxplot comparing QUIESCENCE gene signature scores across tumor GeoMx MRs in Domains A–E and progressing tumors (MEL107–MEL109). Each dot represents a tumor MR (either $SOX10^{+}$ segmented or full tumor MRs). Boxes represent the first and third interquartile of the data; whiskers extend to show the rest of the distribution except for points identified as outliers. **e.** UMAP projection of $SOX10^{+}$ tumor cell clusters obtained by unsupervised Leiden clustering of the CyCIF data. **f.** Scatter plot illustrating the spatial location of tumor cluster 4, colored by cluster label with background cells shown in grey (left).

Supplementary Figure 8.

a.

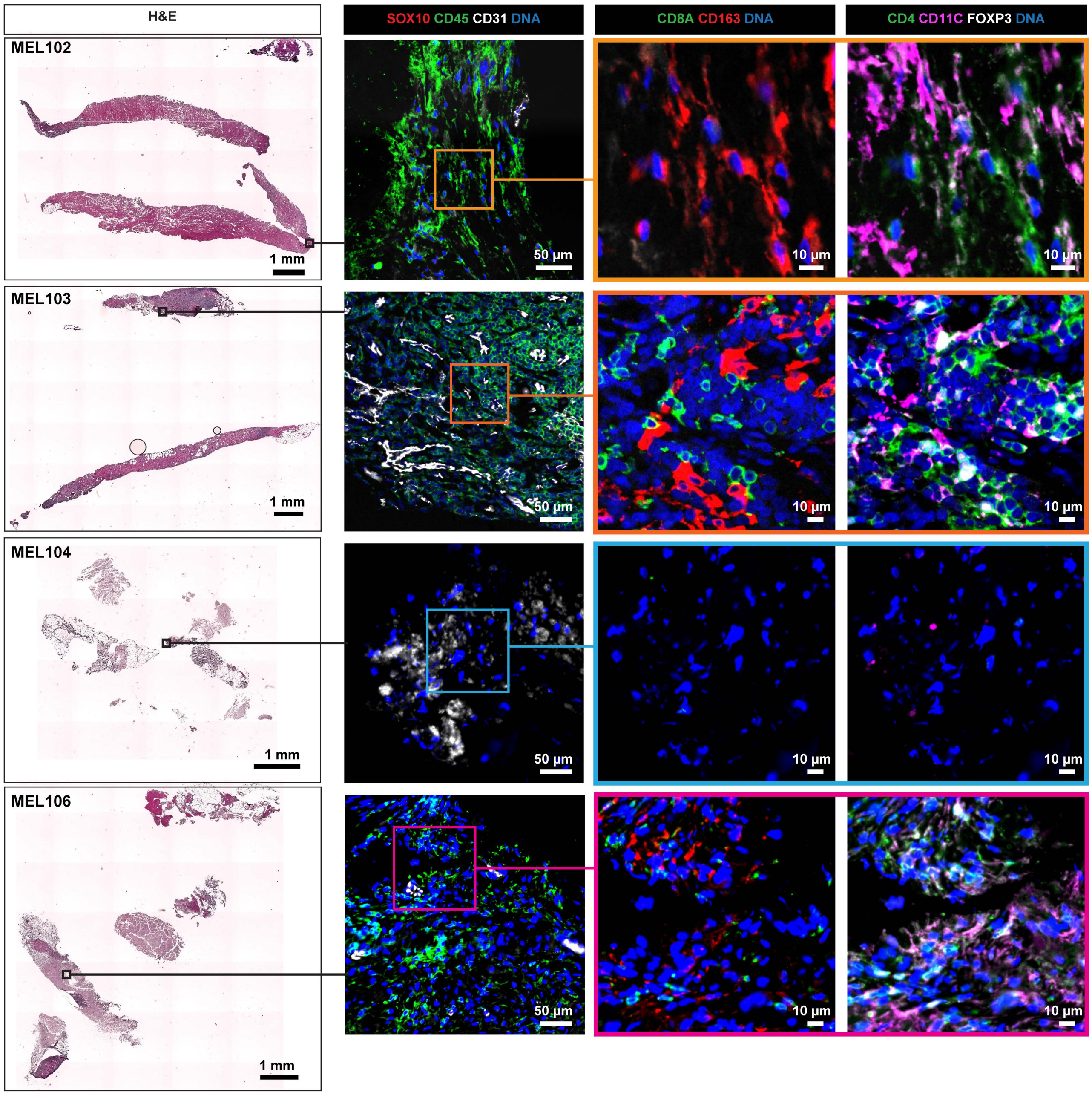


b.

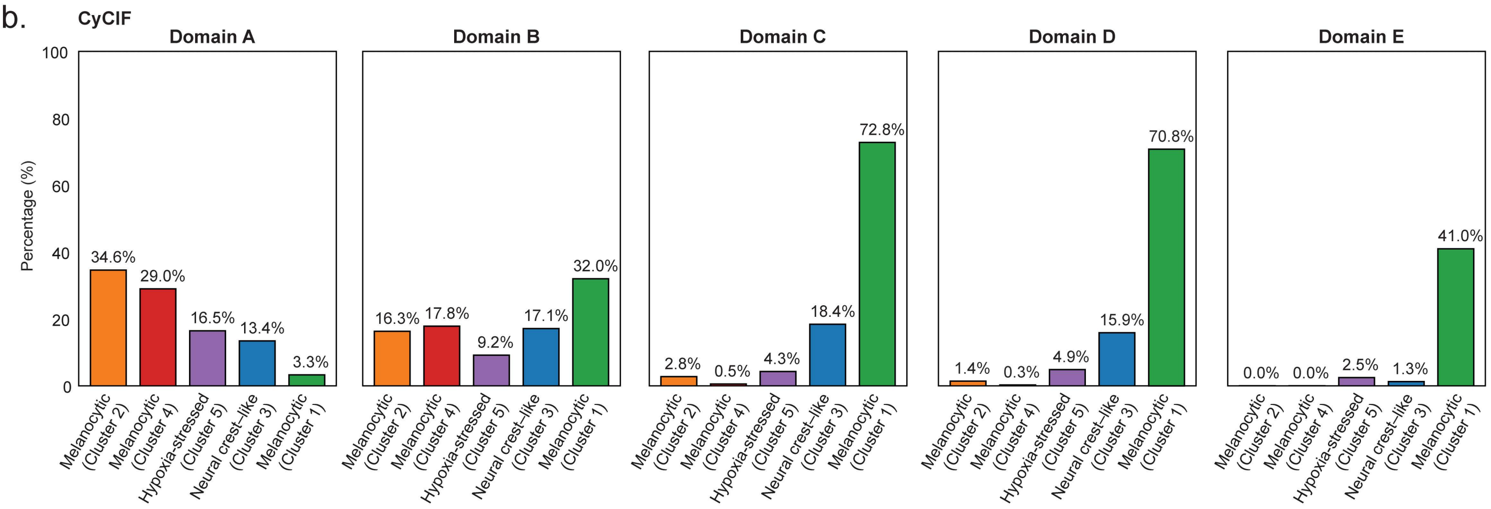

**Supplementary Figure 8. a.** H&E-stained (left) and CyCIF (right) images of specimens MEL102, MEL103, MEL104, and MEL106. CyCIF images were stained for SOX10 (red), CD45 (green), CD31 (white), DNA (blue), CD8A (green), CD163 (red), CD4 (green), CD11C (magenta), and FOXP3 (white). The regions outlined with colored rectangles correspond to magnified regions. Scale bars, 1 mm, 100 μm, and 10 μm. **b.** Bar plots showing the percentage of selected tumor cell clusters defined by CyCIF within each domain. The mean marker expression profiles of each cluster are shown in **Fig. 6c**.
